# Gene capture by transposable elements leads to epigenetic conflict in maize

**DOI:** 10.1101/777037

**Authors:** Aline Muyle, Danelle Seymour, Nikos Darzentas, Elias Primetis, Brandon S. Gaut, Alexandros Bousios

## Abstract

Plant transposable elements (TEs) regularly capture fragments of genes. When the host silences these TEs, siRNAs homologous to the captured regions may also target the genes. This epigenetic cross-talk establishes an intragenomic conflict: silencing the TEs has the cost of silencing the genes. If genes are important, however, natural selection may maintain function by moderating the silencing response, which may also advantage the TEs. Here, we examined this model by focusing on three TE families in maize: Helitrons, Pack-MULEs and Sirevirus LTR retrotransposons. We documented 1,263 TEs containing exon fragments from 1,629 donor genes. Consistent with epigenetic conflict, donor genes mapped more siRNAs and were more methylated than genes with no evidence of capture. However, these patterns differed between syntelog vs. translocated donor genes. Syntelogs appeared to maintain function, as measured by gene expression, consistent with moderation of silencing for functionally important genes. Epigenetic marks did not spread beyond their captured regions and 24nt cross-talk siRNAs were linked with CHH methylation. Translocated genes, in contrast, bore the signature of silencing by being highly methylated and less expressed. They were also overrepresented among donor genes, suggesting a link between capture and gene movement. The evidence for an advantage to TEs was less obvious. TEs with captured fragments were older, mapped fewer siRNAs and were slightly less methylated than TEs without captured fragments but showed no evidence of increased copy numbers. Altogether, our results demonstrate that TE capture triggers an epigenetic conflict for important genes, but it may lead to pseudogenization for less constrained genes.

## Introduction

Transposable elements (TEs) constitute the majority of plant genomes and are major drivers of both genomic and phenotypic evolution (1). TEs are generally not active under normal conditions. Based mostly on studies in *Arabidopsis thaliana*, it is known that this inactivation is achieved by host epigenetic silencing mechanisms that suppress TE activity both before and after transcription (2, 3). These mechanisms rely on small interfering RNAs (siRNAs) that guide *RNAi* and RNA-directed DNA methylation (RdDM) against homologous sequences at the RNA and DNA level, respectively. RdDM is a feedback loop that initiates and spreads cytosine methylation in the CG, CHG and CHH contexts (H = A, C or T) of TE sequences. Symmetric CG and CHG methylation can be then maintained through cell division independent of RdDM, but asymmetric CHH methylation requires continuous *de novo* siRNA targeting (2, 3). As a result, silenced TEs are usually heavily methylated only in the CG and CHG contexts, with CHH methylation typically at much lower levels. Once methylated, silenced TEs are often associated with a closed heterochromatic state (4), which can influence the function and expression of genes, especially when TEs and genes reside in close proximity. For example, methylated and siRNA-targeted TEs can be associated with altered expression of neighbouring genes (5–7) and, as a result, are subject to stronger purifying selection compared to unsilenced TEs or TEs far from genes (6, 8).

In contrast to the epigenetic effects of TEs near genes, much less is known about epigenetic interactions between TEs and genes over long distances, particularly through the *trans-activity* of siRNAs (9). For siRNAs to mediate long distance interactions, there must be sequence similarity between genes and TEs, so that siRNAs are homologous to both. The requirement of sequence similarity can be satisfied by varied evolutionary scenarios, such as the exaptation of portions of TEs into coding genes (10), but it is especially relevant in the phenomenon of gene capture by TEs. Gene capture has been investigated widely in both animals and plants (11, 12). Within plant genomes, capture has been best characterized for Helitrons and Pack-MULE DNA transposons, which together have captured thousands of gene fragments (12, 13). Capture is common enough that a single TE often contains fragments of multiple host genes from unlinked genomic locations (11, 14). Although it is clear that gene capture is common, the mechanisms remain uncertain. However, several mechanisms have been proposed (15, 16), and evidence suggests that capture can occur through both DNA and RNA-mediated processes (14, 17).

The evolutionary consequences of gene capture are also not well characterized. One potential consequence is that the shuffling and rejoining of coding information within a TE leads to the emergence of a novel gene (11). Although this conjecture has been disputed (18), a substantial proportion of TE-captured gene sequences are expressed (12, 19), a subset of those are translated (12, 20), and a few exhibit signatures of selective constraint (13, 18, 20). Another distinct possibility is that gene capture is a neutral mutational process that has few downstream evolutionary ramifications. Finally, gene capture may establish evolutionary conflicts between TEs and genes. Lisch (21) has argued that gene capture is in a TE’s evolutionary interest, because it blurs the line between host and TE “by combining both transposon and host sequences … to increase the cost of efficiently silencing those transposons”. This argument suggests a model of genomic conflict in which a TE captures a fragment from a gene, and the host mounts an siRNA-mediated response against the TE. Because the siRNAs from the captured fragment within the TE can also target the captured region of the ‘donor’ gene (i.e., the gene from which the fragment has been captured), the host response to the TE can simultaneously act in *trans* against the donor gene.

Under this scenario, transcriptional silencing of the TE may have collateral effects on the donor gene, including targeting by siRNAs that lead to DNA methylation and subsequent silencing (Figure 1a). If the donor gene has an important function, then natural selection is likely to either remove affected individuals from the population or limit potential silencing effects on the gene. The latter creates an intragenomic conflict, whereby the advantage of silencing the TE is balanced by potential damage to donor gene function. Conversely, selection to moderate the host response potentially advantages the TE with the captured gene fragment. Notably, this conflict model makes testable predictions that: *i*) donor genes bear the signature of *trans*-epigenetic effects, including increased siRNA targeting and consequent methylation, *ii*) selection may limit these *trans*-epigenetic effects for important vs. less functionally important genes, and *iii*) TEs benefit from capture via decreased host response.

**Figure 1.**
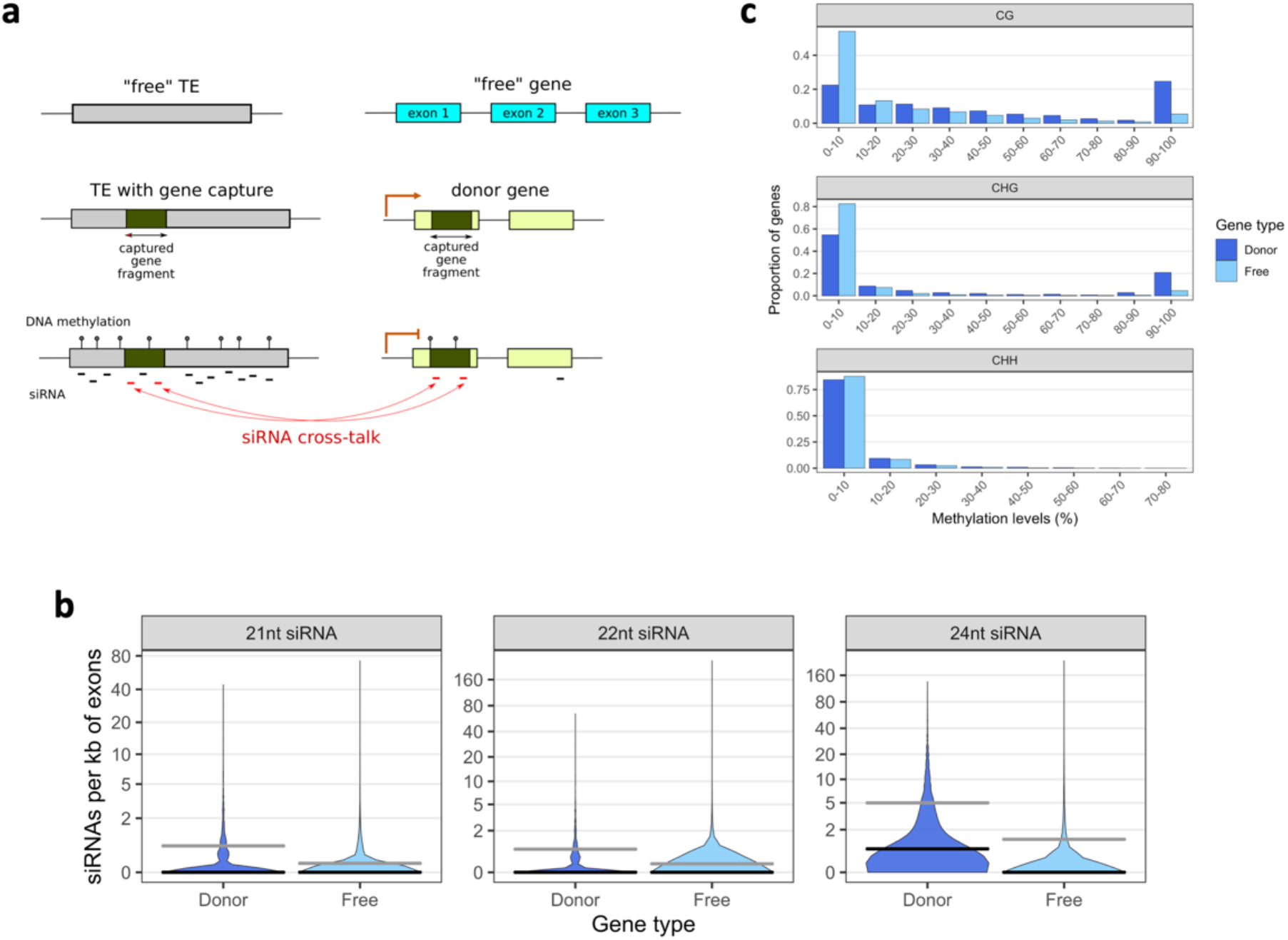
Epigenetic effects of capture on donor genes. **(a)** Schematic of a capture event by a TE and ensuing epigenetic interactions. Definitions used in the text are shown, including donor and free genes, free TEs and TEs with captured fragments, and cross-talk siRNAs that may act in *trans*. The orange arrows indicate expression. **(b)** Number of 21nt, 22nt and 24nt distinct siRNA sequences per kb of exonic mapping to donor and free genes. The gray lines indicate the mean, the black lines the median. **(c)** Distribution of the proportion of CG, CHG and CHH exonic methylation of donor and free genes. Data are from the ear tissue.

The possibility of epigenetic links between TEs and donor genes has been discussed previously (11), but to our knowledge only one study has examined how often siRNAs map to both donor genes and to their captured fragments (20). This study focused on Pack-MULEs in rice (*Oryza sativa*) and found siRNAs that map to both TEs and donor genes, thus providing the potential for siRNA ‘cross-talk’ between donor genes and captured gene fragments. The study also found that genes with cross-talk are less expressed compared to genes without any mapped siRNAs. Two recent studies of rice Pack-MULEs extended this line of enquiry by investigating whether donor genes are methylated (12, 19), which could be indicative of epigenetic effects consistent with the conflict model. They found, however, that donor genes have low methylation levels that do not differ substantially from genes with no apparent history of capture by TEs (hereafter termed ‘free’ genes). These studies provide some, but limited, evidence for epigenetic conflict.

The study of Pack-MULEs in rice suffers from two potential shortcomings with respect to investigating epigenetic interactions. The first is Pack-MULEs themselves. They commonly capture genes and therefore provide a rich template for study, but often have lower methylation levels than other TE families (12, 22), possibly because they preferentially insert near the 5’ termini of genes (23). This tendency may lessen the potential for intragenomic conflict with their donor genes. The second shortcoming is the small genome size of rice. Large genomes differ from small genomes in their TE content and, as a result, their DNA methylation patterns. For example, Takuno et al. (2016) showed that only 6% of genes in rice (490 Mb) and 2% of genes in *A. thaliana* (156 Mb) have high levels of methylation (≥90% of methylated cytosines) in the CG context compared to 24% of genes in the much larger (2,646 Mb) genome of maize (24). The contrast is even stronger for the CHG context where 12%, 1% and <1% of maize, rice and *A. thaliana* genes, respectively, have high methylation levels (≥90%), reflecting the strong positive correlation between gene CHG methylation and genome size (24–26). As a result of these differences, large genomes like maize may represent better systems to study gene capture by TEs and the impact on donor genes.

Here, we hypothesize that gene capture may have epigenetic consequences for endogenous genes in maize. To test this hypothesis, we identify capture events representing all three major TE classes, i.e. Helitron rolling circle transposons, Pack-MULE Class II DNA transposons, and a representative of Class I retroelements, Sirevirus LTR retrotransposons (27). Sireviruses are crucial because they compose ~20% of the maize genome (28), are targeted by large numbers of siRNAs, and are highly methylated (29). Given sets of TEs with gene capture events, we integrate evolutionary analyses with siRNA, methylation and gene expression data to address two sets of predictions. The first set focuses on the genic viewpoint. If the conflict model holds, we predict that donor genes bear the signature of *trans*-epigenetic effects compared to free genes. We also predict that natural selection will moderate these epigenetic effects on functionally important genes relative to less important genes. In the second set of predictions, we focus on TEs with captured gene fragments. Is there any evidence that they benefit from gene capture via decreased host response?

## Results

### Identifying captured gene fragments and their donor genes

To investigate the potential for intragenomic conflict, we first identified gene capture events. Identifying true events is a challenging task, because annotation errors can lead to false positives that mislead downstream analyses. Annotation errors can be particularly pernicious for TEs, because a proportion of putatively full-length elements may represent partial sequences or mosaics of different TEs. This ambiguity is evident in TE annotations of the recent B73 RefGen_v4 genome that predict, for example, different numbers of Helitron sequences, 49,235 (30) vs. 22,339 (22), even though both used HelitronScanner (31). To address this concern, we favored specificity over sensitivity by using previously published and carefully curated smaller datasets of full-length elements for Helitrons (31), Pack-MULEs (23), and Sireviruses (32). These datasets were mostly based on RefGen_v2 and contained 1,351, 275 and 13,833 elements respectively. For example, the 1,351 Helitrons represented a high-quality subset of 31,233 full-length elements identified in the original HelitronScanner manuscript that were, however, additionally validated in the same study with *in silico* comparisons with the genome of the Mo17 inbred line (31). We curated these datasets to further remove problematic elements and converted their chromosomal coordinates to RefGen_v4 to ensure that these TEs are physically present in the most recent genome version (see Methods). Overall, we generated a sample of 7,473 TEs, which consisted of 1,035 Helitrons, 238 Pack-MULEs and 6,200 Sireviruses. We implemented a similarly strict pipeline for the 39,423 genes of the Filtered Gene Set (FGS) to remove low-quality candidates (e.g. possible misannotated TEs) and 5,495 genes that were no longer annotated in RefGen_v4 (see Methods). The final dataset consisted of 27,056 genes.

We then performed strict BLASTN comparisons (*E*-value cutoff of 1×10^-40^) between the TEs and the exons of the genes to identify both captured gene fragments within TEs and their donor genes. We only kept hits that belonged to the longest alternative transcript of each gene, removed cases of physical overlaps between full-length TEs and complete genes, and used the BLASTN bit score to select the true donor gene when exons from multiple candidates generated overlapping hits within a TE (20, 23). This approach derived a final set of 1,629 donor genes (out of 4,117 candidate genes with hits to TEs), with the remaining 22,939 genes characterized as free genes. Several features of the donor genes suggest that they are not pseudogenes or small gene fragments located within TEs. For example, similar proportions of donor and free genes were assigned a specific function in RefGen_v4 (82.2% vs. 84.4%), and donor genes had more and longer exons than free genes (Figure S1). The donor genes were captured by 1,263 distinct TEs. Most Helitrons (873; 84%) and Pack-MULEs (186; 78%) contained gene fragments, in contrast to a much smaller proportion of Sireviruses (204; 3%). Like previous studies (11, 14), we found that individual elements often contained multiple independent capture events: 68% of Helitrons harbored ≥2 captured fragments, as did 50% of Pack-MULEs and 15% of Sireviruses. Finally, we confirmed that the elements of each family had sequence or structural features that correspond to full-length TEs, i.e. the conserved 5’ and 3’ termini for Helitrons (31), terminal inverted repeats for Pack-MULEs, and short target-site duplications for Sireviruses (Figure S2).

### Donor genes are targets of siRNAs and are highly methylated

Under our conflict model, the first prediction is that gene capture should lead to siRNA cross-talk between genes and TEs, potentially leading to increased methylation of donor genes. Accordingly, we contrasted the exons of donor and free genes for siRNA mapping and methylation characteristics. Throughout this study, we relied on published siRNA and bisulfite-sequencing (BS-seq) datasets, focusing on libraries from unfertilized ears (33, 34), leaves of maize seedlings (35, 36) and tassels (37) (see Methods). We analyzed 21nt, 22nt, and 24nt siRNAs (both uniquely and multiply mapped in the genome), because these lengths are involved in TE silencing. Considering, however, that 21nt/22nt siRNAs mostly participate in *RNAi*/post-transcriptional silencing while 24nt siRNAs in RdDM/transcriptional silencing (2, 3), we analyzed each length separately. For each gene, we calculated the number of distinct siRNA sequences per kb of all their exons combined, a metric that avoids the errors inherent to measuring siRNA expression (38). Across all genes and libraries, exonic mapping was strongly correlated for 21nt vs. 22nt siRNAs (mean Pearson coefficient r=0.85, p=0) but not for 21nt/22nt vs. 24nt siRNAs (mean Pearson coefficient r=0.58, p=0), likely reflecting their roles in different epigenetic pathways. Results were generally consistent among tissues; hence, we report data from ear in the main text, and provide results from the other two tissues mostly in Supplementary Information.

The comparison of the siRNA mapping profiles of the exons of donor and free genes revealed striking differences: the 1,629 donor genes mapped more siRNAs per kb than the 22,939 free genes (Figure 1b, S3, one-sided Mann-Whitney U test p<2.2e-16 for all siRNA lengths and tissues). Across all tissues combined, donor genes mapped 3.0 times more 24nt siRNAs per kb on average than free genes, compared to 1.7 times for 21nt and 22nt siRNAs respectively. Differences in siRNA mapping are expected to affect methylation patterns. We calculated the proportion of methylated cytosines in the CG, CHG and CHH contexts of exons using only uniquely mapped BS-seq reads across the genome. Of the 1,629 donor and 22,939 free genes, 1,525 and 21,614 passed CG methylation filters (≥10 covered CG sites), representing ~94% of the genic dataset, with similar proportions retained for CHG and CHH methylation. We found that the distribution of CG and CHG methylation was notably bimodal, with most genes having either low (≤ 10%) or high (≥90%) methylation (Figure 1c). This pattern is consistent with previous work on several plant species (24, 25). However, donor and free genes generated strikingly different distributions, showing a bias towards high and low methylation, respectively: in ear, 24.9% of donor genes had ≥90% of their cytosines methylated in the CG context, and 20.9% in the CHG context. In contrast, only 5.5% of free genes were highly methylated in the CG context, with 4.6% in the CHG context. In fact, most free genes had ≤ 10% CG and CHG methylation, 54.1% and 82.6% respectively (Figure 1c). As expected, methylation in the CHH context was much lower, because the majority of genes in both datasets had low (≤5%) levels of methylation. However, although not clearly evident in the histogram, donor genes were significantly more methylated than free genes, with a mean of 5% vs. 3.7% (one-sided Mann-Whitney U test, p<2.2e-16), and a higher proportion with high (≥15%) CHH methylation - i.e. 10.6% vs. 6.6% of free genes. Overall, the trends were clear and consistent across all tissues (Figures 1, S3, S4): donor genes mapped more siRNAs and were more highly methylated.

### Dramatic differences in the epigenetic profiles of syntenic vs. non-syntenic donor genes

Our results support the predictions of the conflict model by showing that donor genes are heavily enriched for both siRNA mapping and methylation levels. However, the model specifically proposes that intragenomic conflict arises for functional genes, but many donor genes have high levels of methylation, especially in the CHG context (Figure 1c), which is a potential signature of silencing. To better test the conflict model, we split the genic dataset according to syntenic relationships with *Sorghum bicolor* (39). Our reasoning was that syntenic orthologs are enriched for genes that are functional and associated with phenotypes (40). We thus expect that these genes are more often subject to selective constraint and hence susceptible to epigenetic conflict. In contrast, non-syntenic genes are more likely to be dispensable or non-functional (40) and thus more likely to escape selective constraint. We therefore predict that, as a general trend, the conflict model should not hold for non-syntenic genes.

We assigned the 1,629 donor and 22,939 free genes into two categories: syntenic orthologs (hereafter ‘syntelogs’) and genes that have moved their location in maize relative to sorghum (hereafter ‘translocated’) (see Methods). The two categories yielded a striking observation: translocated genes had a higher probability than syntelogs to be captured by TEs. In total, 58.2% (948) of donor genes were syntelogs and 27.1% (442) were translocated, while their proportions among free genes were 78.7% (18,046) and 9.8% (2,243) respectively (Chi-squared=512.37, p<2.2e-16). We also note that a much higher proportion of donor and free syntelog genes (91.6% and 87.2%) was assigned a specific function compared to translocated genes (64% and 63.3%) based on the RefGen_v4 annotation, which supports our contention that syntelogs are more likely to be functionally important. Most of the remaining donor (201; 12.3%) and free (2,284; 10%) genes were either located in regions of maize chromosomes that were not identified in sorghum or completely lacked synteny information. We excluded these two categories from further analyses due to their ambiguous syntenic status.

We then contrasted the epigenetic profiles of syntelog and translocated genes, starting with siRNA mapping. Looking *within* each synteny-based category first, differences between donor and free genes remained consistent to our analysis based on all genes, i.e, donor genes were targeted by more siRNAs per kb than free genes across all tissues (Figure 2a, S5a, one-sided Mann-Whitney U test p<2.2e-16 for most combinations). This result was largely due to the higher fraction of free genes that did not map any siRNAs, with this difference being more prominent for 24nt siRNAs, where twice as many free genes had no mapping events compared to donor genes (Figure 2b). Removing genes with no siRNAs did not change the mapping differences within syntelogs but did so for translocated genes because donor and free genes were equally targeted by siRNAs (Figure S5b). For syntelog genes and across all tissues combined, donor genes mapped 3.8-fold more 24nt siRNAs per kb on average than free genes, compared to 2.1-fold for both 21nt and 22nt siRNAs. These differences in average siRNA mapping were not as strong within translocated genes; for example, donor genes mapped 1.4-fold more 24nt siRNAs than free genes.

**Figure 2.**
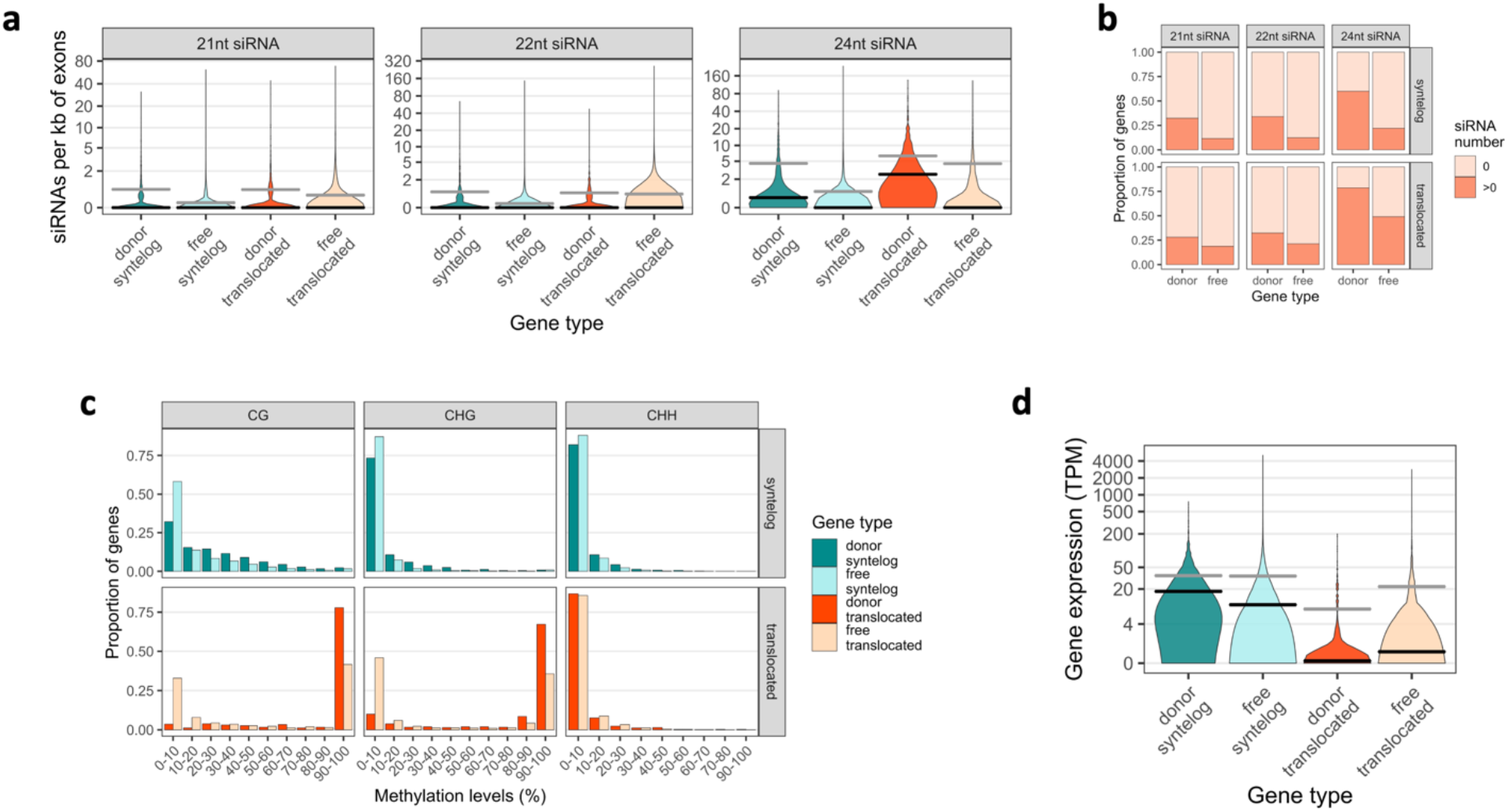
Epigenetic and expression profiles of donor and free genes split by their syntenic status with sorghum. The four gene categories in all plots are donor syntelogs, free syntelogs, donor translocated and free translocated genes. **(a)** Number of 21nt, 22nt and 24nt distinct siRNA sequences per kb of exonic mapping. **(b)** Proportion of genes with no siRNA mapping. **(c)** Distribution of the proportion of CG, CHG and CHH exonic methylation. **(d)** Gene expression measured in TPM. Data are from the ear tissue. The gray lines in panels (a) and (d) indicate the mean, the black lines the median.

Noting that donor genes had higher levels of siRNA targeting than free genes within each synteny-based category, the differences *between* categories were intriguing. Across libraries, the most striking pattern was that the level of 24nt siRNA targeting of donor translocated genes was higher than any other category (Figure 2a, S5a; one-sided Mann-Whitney U test p<2.2e-16). This was not the case for 21-22nt siRNAs that often did not map at levels statistically different between syntelog and translocated genes (Figure 2a, S5a). Taken together, these results lead to three main observations. First, gene capture by TEs is linked to increased levels of siRNA targeting in donor genes (e.g. donor syntelog vs. free syntelog genes); second, these levels are lower in ‘important’ compared to ‘less important’ genes (e.g. donor syntelogs vs. donor translocated genes); and third, 24nt siRNAs appear to be the crucial component of these differences.

The corresponding methylation patterns of the four gene categories supported the siRNA results. The distribution of CG and CHG methylation in donor and free syntelogs was no longer bimodal due to the absence of genes with high methylation (Figure 2c, S6). However, donor syntelogs still had higher methylation levels than free syntelogs (in ear mean CG 26.7% vs. 15.5%, one-sided Mann-Whitney U test p<2.2e-16; CHG 9.6% vs. 4.9%, p<2.2e-16), including in the CHH context (mean 5.3% vs. 3.6%, p=3.5e-11), where twice as many donor syntelogs had high methylation (12.6% vs. 6.5%). The methylation profile of translocated genes contradicted the above patterns (Figure 2c, S6). The majority of translocated donor genes had high CG (76.7%) and CHG (66.8%) methylation and virtually none had low methylation, while translocated free genes were clearly distinguished by having a balanced bimodal distribution of methylation (Figure 2c, S6). Unlike syntelogs, CHH methylation was more similar between the two sets of translocated genes (in ear mean 4.9% vs. 4.6%, p=0.0052; 7.8% of donor vs. 8.4% of free with high CHH methylation). Overall, the methylation patterns recapitulated the siRNA patterns by suggesting that donor genes tended to be more methylated than free genes, which in the case of donor translocated genes reached very high levels.

### Captured regions of syntelog donor genes are enriched for repressive epigenetic marks

An additional prediction of the conflict model suggests that the epigenetic marks of siRNA mapping and methylation should be overrepresented in the regions that were captured by TEs, at least for functionally important genes (Figure 1a). To examine this prediction, we compared siRNA mapping and methylation levels between the captured vs. non-captured exonic regions of donor genes. As predicted, significantly more siRNAs mapped to the captured than the non-captured regions of syntelogs, with 24nt siRNAs generating the strongest difference across libraries (one-sided Wilcoxon signed rank test p<2.2e-16; Figure 3a, S7a). The captured regions of syntelogs were also significantly more methylated in the CG (in ear one-sided Wilcoxon signed rank test p=9.45e-12), CHG (p=4.36e-12), and CHH (p=1.49e-05) contexts (Figure 3b, S7b). Supporting the statistical tests, high levels of methylation were found only in captured regions of syntelogs; in ear, 27.3% of captured regions had ≥90% of their CG sites methylated vs. only 1.4% for non-captured regions. These differences extended to high CHG (15.6% captured vs. 0.2% non-captured regions) and high CHH (22% captured vs. 6.5% non-captured regions) methylation. This result was robust when we increased the coverage filter in each locus from ≥10 to ≥40 covered sites, which tended to exclude captured regions with short lengths that could theoretically bias the results (Table S1). In contrast to syntelogs, the captured and non-captured regions of translocated genes did not significantly differ in either siRNA targeting or methylation levels in most cases (Figure 3, S7). Altogether, these results suggest that repressive epigenetic marks may not spread beyond the captured regions of important syntelog genes but do so in translocated genes.

**Figure 3.**
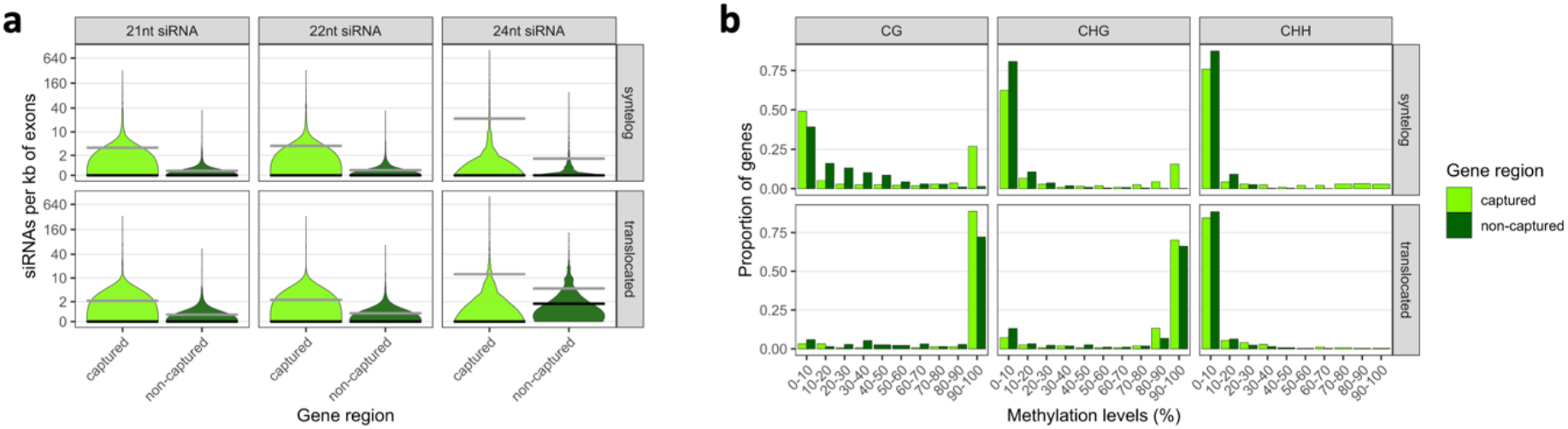
siRNA and methylation patterns of captured and non-captured regions of syntelog and translocated donor genes. **(a)** Number of 21nt, 22nt and 24nt distinct siRNA sequences per kb of captured and non-captured exonic regions. The gray lines indicate the mean, the black lines the median. **(b)** Distribution of the proportion of CG, CHG and CHH exonic methylation of captured and non-captured regions. Data are from the ear tissue.

### 24nt cross-talk siRNAs are enriched in donor syntelog genes and affect CHH methylation in trans

Our conflict model is based on cross-talk siRNAs, which are siRNAs that map both to the captured fragment within the TE and the gene and may act *in trans*. To test whether cross-talk siRNAs represent an enriched fraction of the total number of siRNAs that mapped to donor genes, we employed a binomial test that compared the observed proportion of cross-talk siRNAs (cross-talk siRNAs / all siRNAs) to the proportion of captured gene length (captured exon length / total exon length) across all genes. In each tissue, the binomial test revealed a significant enrichment of cross-talk siRNAs in donor syntelogs (Table S2), and this was especially strong for 24nt siRNAs (p~0). Translocated genes had significantly fewer cross-talk siRNAs in leaf and tassel, and moderate statistical support for enriched cross-talk in ear (Table S2). These contrasting patterns suggest that syntelogs are disproportionately targeted by cross-talk siRNAs, while translocated genes are targeted more generally.

A key prediction of the conflict model - i.e. that gene capture has the capacity to modify the epigenetic state of the donor gene - presupposes that cross-talk siRNAs can act in *trans*. Hence, we used a linear model with mixed effects across all tissues (see Methods) to examine the relationship between cross-talk siRNAs and methylation of captured and non-captured regions of donor genes. We found that the number of 24nt cross-talk siRNAs was positively correlated to the methylation levels of captured regions within donor syntelogs (Table 1). This was especially true for CHH methylation, where 21.5% of the variance across captured regions was explained by the abundance of 24nt crosstalk siRNAs, but 10.7% of the variance was also explained for CHG methylation and 1.6% for CG methylation. The fact that more variation was explained for CHH methylation makes biological sense, because methylation in this context is maintained *de novo* by 24nt siRNAs via RdDM (3). In contrast, the methylation levels of the non-captured regions were not positively associated with the abundance of 24nt cross-talk siRNAs (Table 1). In addition, these patterns did not hold as clearly for the other siRNA lengths for donor syntelogs or, generally, for translocated genes (Table S3). For example, only 2.6% and 4.7% of the variance of CHH methylation within the captured regions of donor syntelogs was explained by 21nt and 22nt cross-talk siRNAs respectively. Taken together, these results establish an epigenetic link in donor syntelogs between gene capture, 24nt cross-talk siRNAs that act *in trans*, and methylation, particularly in the CHH context.

**Table 1.** Correlation between the number of 24nt cross-talk siRNAs and methylation levels of captured and non-captured regions of donor syntelog genes

| region | methylation context | Estimate | Standard Error | t-value | p-value | marginal R<sup>2</sup> |
| --- | --- | --- | --- | --- | --- | --- |
| captured | CG | 2.82e-3 | 5.41e-4 | 5.21 | 2.21e-7 | 0.0156 |
|  | CHG | 0.00577 | 0.000455 | 12.7 | <2e-16 | 0.107 |
|  | CHH | 0.00513 | 0.000280 | 18.3 | <2e-16 | 0.215 |
| non-captured | CG | -0.001326 | 0.0003339 | -3.97 | 0.0000746 | 0.0096 |
|  | CHG | no significant effect (p=0.1402) |  |  |  |  |
|  | CHH | no significant effect (p=0.5697) |  |  |  |  |

The fact that siRNA cross-talk is significant for syntelog genes raises an interesting question: what is the relationship between siRNA cross-talk and the time since the capturing event took place? This is probably a complex relationship, for two reasons. First, the initiation of the host epigenetic response against a new capture event may not be immediate, so that very recent capture events may not generate enough siRNAs to detect cross-talk. Second, the opportunities for cross-talk are finite, because the sequences of the donor gene and the captured fragment within the TE diverge over time. As they diverge, cross-talk can no longer occur as efficient because siRNAs no longer match both entities. We used synonymous divergence (*d_s_*) between the donor syntelog and the TE-captured exon as a proxy of the age of capture. We then examined the relationship between the abundance of siRNA cross-talk and time since gene capture. The tests were significant and positively correlated only when we combined all siRNA lengths. This correlation suggests that donor syntelogs with older capture events had more cross-talk siRNAs over time, despite the increased divergence of their captured sequences (linear model with mixed effects across all tissues z-value=2.06, *p*=0.0398, marginal R-squared 0.006, see Methods). Overall, we interpret these results to imply that it takes time for cross-talk to evolve after the capture event.

### Capture does not affect the expression of donor syntelog genes

Our analyses are consistent with the interpretation that cross-talk siRNAs drive, to some extent, methylation of donor genes. The conflict model predicts, however, that these epigenetic modifications will have minimal effects on important genes, because natural selection acts against changes that affect function. To test this conjecture, we contrasted expression patterns of donor and free genes using data from the Atlas Expression database (see Methods). Indeed, we did not find significantly lower levels of expression in donor syntelog compared to free syntelog genes in ear, leaf and ten different cell types of the maize kernel (Figure 2d, S8). In fact, donor syntelogs were expressed at significantly higher levels (log transformed average of Transcripts per Million, TPM, 1.88 vs. 1.23 across tissues, onesided Mann-Whitney U test p between 2.7e-08 and 2.2e-16) and had a lower proportion of genes with no expression (average of 11.5% vs. 24% across tissues) than free syntelogs. In contrast, both categories of translocated genes had lower levels of expression compared to syntelog genes (Figure 2d, S8), which is in agreement with previous studies that showed translocated genes to have pseudogene-like characteristics (40, 41). Donor translocated genes, however, appeared to be driving this difference because they had significantly lower expression than free translocated genes (log transformed average TPM across tissues −0.52 vs. 0.32, one-sided Mann-Whitney U test p between 0.0009 and 8.23e-12). Therefore, donor translocated genes exhibit a signal consistent with run-away epigenetic interactions with TEs that are not moderated by functional constraints, hence dramatically reducing expression.

Finally, we examined if gene expression is affected by the position of the captured fragment within the gene. For example, it is possible that capture and subsequent epigenetic interactions at the 5’ or 3’ untranslated regions (UTRs) may affect expression levels, because these are regions of major importance for gene regulation (42, 43). To investigate this, we classified each gene based on whether the captured fragment(s) were part of the 5’ or 3’ exons to approximate the location of UTRs, or any internal exon. By analyzing genes whose captured fragment(s) were derived from a single position only, we found that capture of the 5’ or 3’ exons of syntelogs significantly reduced expression across all tissues compared to capture of internal exons (one-sided Mann-Whitney U test p between 1.9e-08 and 2.2e-16), while this was not the case for translocated genes (Figure S9). We note, nevertheless, that the expression levels of these donor syntelogs were still higher than free syntelogs (one-sided Mann-Whitney U test p between 0.02951 and 2.1e-06 across tissues).

### Potential advantages for TEs to capture gene fragments

Besides the impact on genes, the conflict model also predicts that TEs with captured gene fragments gain an advantage, due to a moderation of the host response. To explore this possibility, we focused on 852 TEs that captured at least one fragment from a syntelog and contrasted them to 5,931 ‘free’ TEs that had no BLASTN hit to the gene dataset. Given these two groups, we considered four potential measures of advantage for TEs with syntelog capture: *i*) they may be retained within the genome for longer lengths of time, *ii*) they may be targeted by fewer siRNAs, *iii*) they may have lower levels of methylation, and *iv*) they may proliferate more often, leading to higher copy numbers.

To test the first idea, we used age estimates from terminal branch lengths of TE phylogenetic trees generated by Stitzer at al. (2019). We found that TEs with syntelog capture are older than free TEs (mean of 0.135 vs. 0.066 million years, one-sided Mann-Whitney U test p<2.2e-16; Figure 4a), suggesting that they have remained intact within the genome for longer periods. They also mapped significantly less siRNAs of all lengths based on a linear model across all tissues and after removing the captured regions from TEs with captured fragments (for example, contrast for 24nt siRNAs z-ratio=-57.59, p<0.0001, marginal R-squared=15.55%, see Methods) (Figure 4b, Figure S10a). This result remained significant after including TE age in the model (Table S4). We also found that TEs with syntelog capture were less methylated than free TEs in both the CG (ear mean 95.5% vs. 98.5%) and CHG (89.4% vs. 91.4%) contexts (Figure 4c, Figure S10b). These differences were small but significant for CG methylation in a linear model across all tissues and held after controlling for TE age (CG: contrast z-value=4.71, p=2.44e-06; CHG contrast p=0.695; Table S5). However, TEs with syntelog capture had significantly more CHH methylation compared to free TEs (12.0% vs. 3.1%, contrast t-value=-25.86, p<2e-16, Figure 4c, Figure S10b, Table S5). Finally, to address the issue of copy number, we assessed how many times a donor syntelog was found within multiple TEs and found that the majority (71.2%) were in a single element, and this proportion was not statistically different from donor translocated genes (67.2%) (Figure 4d). This suggests that TEs do not amplify in large numbers following syntelog gene capture.

**Figure 4.**
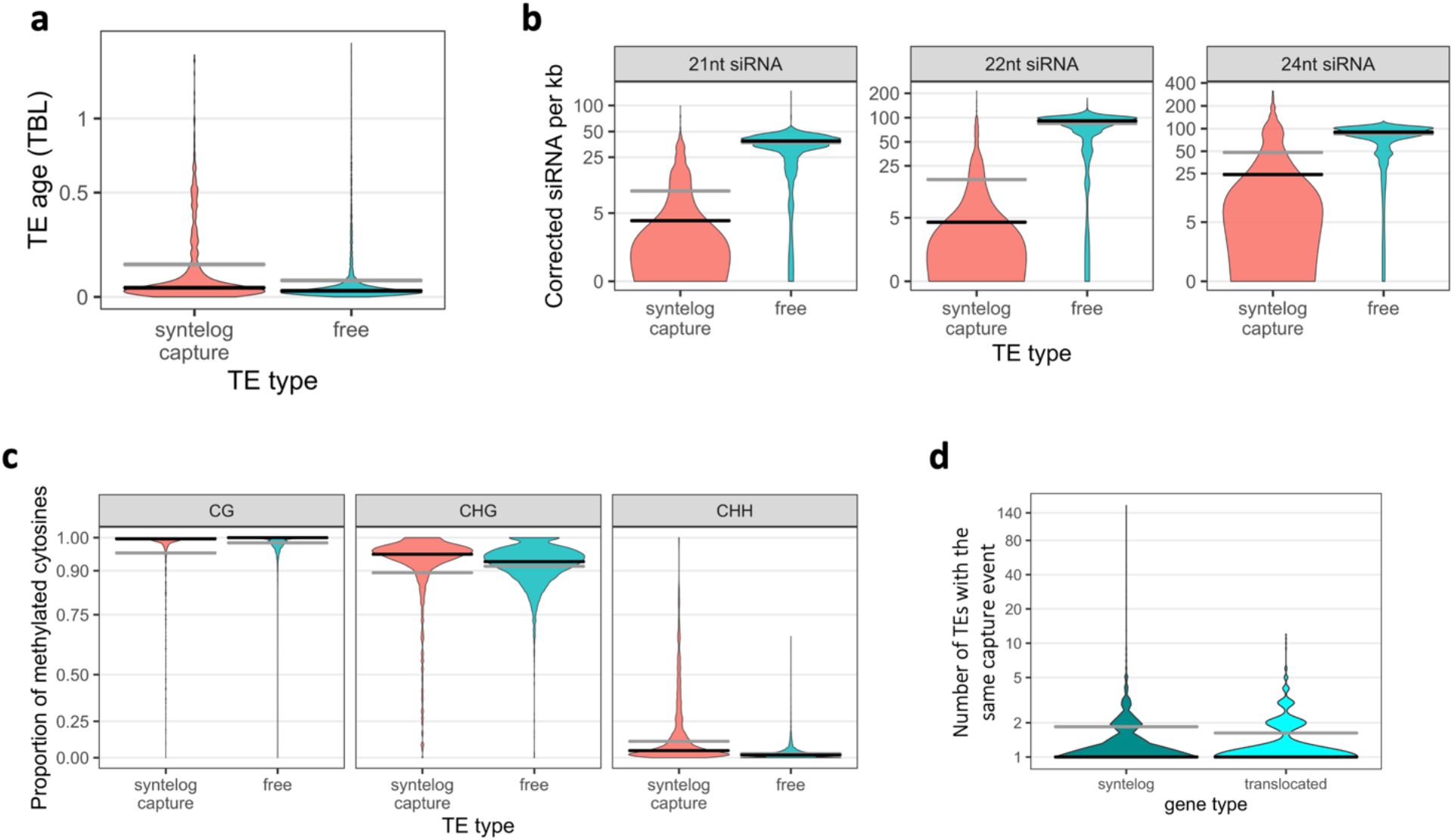
Characteristics of TEs with syntelog capture vs. free TEs. **(a)** TE transposition age in terminal branch lengths (TBL). **(b)** Number of 21nt, 22nt and 24nt distinct siRNA sequences per kb mapping to TEs. This was computed after removing captured regions from TEs, but results were qualitatively identical when they were included. **(c)** Proportion of methylated cytosines in CG, CHG and CHH contexts of TEs. **(d)** Distribution of the number of TEs with the same capture event for TEs that have captured syntelog vs. translocated genes. The gray lines indicate the mean, the black lines the median.

### Patterns of conflict are reproducible at the TE family level

Thus far we have primarily reported analyses based on all three TE families together. We combined families to provide a large number of observations for both syntelog and translocated genes, but, importantly, to also provide a global view of the epigenetic effect of TE capture on host genes. We did, however, examine each family separately and found that the main results were reproducible at the family level (summarized in Table S6, S7): i) donor genes mapped more siRNAs and were more methylated than free genes in both syntenic categories; ii) the hotspot for these epigenetic patterns was the captured region for donor syntelogs; iii) the expression level was higher for donor syntelogs compared to free syntelogs and, conversely, it was lower for donor translocated vs. free translocated genes. We note, however, that the expression of genes captured by Pack-MULEs was not statistically different to free genes either for syntelogs or translocated genes. Finally, we repeated the analysis for TE advantage, and only Sireviruses generated significant trends but only for age (mean of 0.0972 vs. 0.0652 million years, one-sided Mann-Whitney U test p=6.14e-06) and siRNA mapping (Table S4 and S5). The lack of significance for Helitrons and Pack-MULEs may reflect the fact that few of these elements lack captured gene fragments.

Overall, these findings suggest that gene capture by any TE family is likely to trigger similar downstream epigenetic effects. However, we also generated evidence that TE families may be capturing genes in distinct ways. For example, Helitrons and, to a lesser extent, Sireviruses exhibited a preference for translocated genes: compared to their proportion of 9.8% among free genes, translocated genes represented 30.4% (412/1,356) of the genes captured by Helitrons (Chi-squared=556.34, p<2.2e-16) and 20.3% (30/148) of those captured by Sireviruses (Chi-squared=17.075, p<1.8e-05) (Figure S11a). In contrast, only 7.5% (15/200) of the Pack-MULE genes were translocated. Furthermore, the three families tend to capture different parts of genes. Helitrons exhibited a preference for internal exons (45.3%), but Pack-MULEs most often captured 5’ exons (42.1%) and Sireviruses 3’ exons (64.2%) (Figure S11b). If the capture of internal exons has smaller effects on donor gene expression (Figure S9), then Helitrons may cause less functionally impactful epigenetic conflicts than the other families. The orientation of the captured fragments also differed among families. While there was no orientation bias for Pack-MULEs and Sireviruses, 78.6% of captured fragments of Helitrons were in the sense orientation (Figure S11c). If antisense capture events can trigger *RNAi* more readily during TE expression, then their deficit in Helitrons may again limit some aspects of their epigenetic effects on donor genes.

## Discussion

TEs are often in conflict with their plant hosts, because their proliferation tends to have a deleterious effect on host fitness. While this aspect of the TE-host conflict is well established, here we have studied a unique aspect of their conflict, which is driven by gene capture and ensuing epigenetic interactions between TEs and genes. To study this conflict, we have formalized a model suggested by Lisch (21). He argued that gene capture can have a beneficial effect on TEs because they become ‘camouflaged’ and, hence, are less apt to be silenced by the host epigenetic machinery. We have extended the model to also consider the effect of capture on donor genes, predicting that they should have higher siRNA mapping relative to genes with no history of capture. Moreover, if the TE is subjected to silencing, we predict that siRNA cross-talk between the TE and the gene drives epigenetic alterations to the gene itself. The epigenetic modification of the donor gene may eventually reach a threshold that affects gene function, ultimately driving intragenomic conflict, especially if the gene is functionally important. That is, when the silencing response against the TE becomes deleterious to the donor gene, then natural selection may favor an amelioration of the silencing response (Figure 5a).

**Figure 5.**
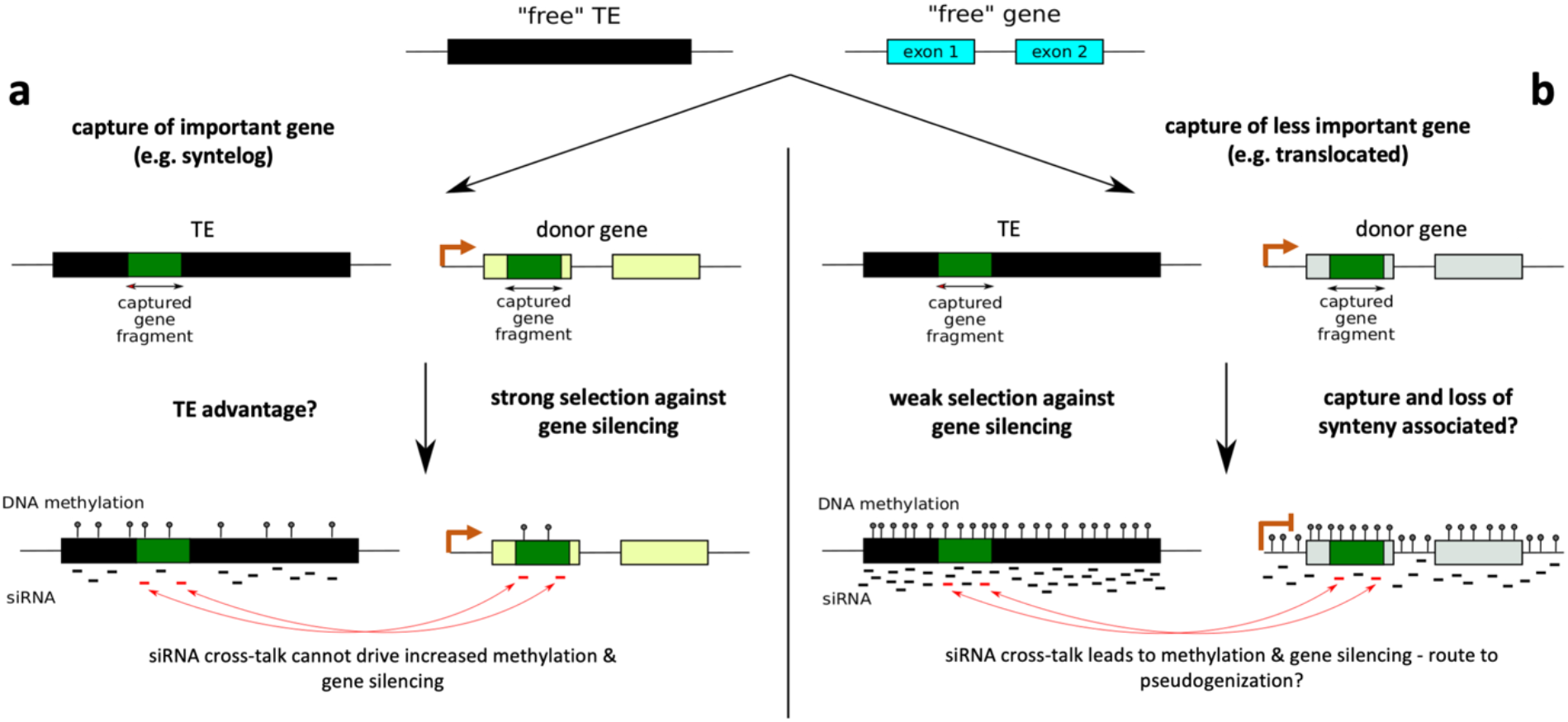
The epigenetic conflict model of gene capture. When TEs capture fragments of genes, siRNAs derived by the TEs may act *in trans* to accidentally mediate an epigenetic response against the gene, leading to increased methylation and reduced expression. The conflict comes from evolutionary pressure to silence TEs without simultaneously silencing functionally important genes, syntelogs in our example. As a result, epigenetic effects on these genes are moderated by natural selection and expression is not affected. TEs may benefit from this moderation, although this remains unclear. In contrast, for genes that are not under strong selective constraint, methylation can increase in the absence of conflict, leading to loss of expression and potential pseudogenization. This profile is characteristic of genes that have moved from their syntenic loci, which are overrepresented among donor genes, suggesting that capture may trigger movement.

### The case for conflict: genes

What is the evidence to support this model? Based on our dataset of syntelogs - to which the conflict model should apply because they are enriched for functionally important genes according to both our analysis and previous studies (40) - we find that donor genes map more siRNAs and are more highly methylated than free genes (Figure 2a, 2c). These epigenetic markers are enriched in the captured regions of genes (Figure 3), where a large fraction of siRNAs also map (cross-talk) to the captured fragment within the TE (Table S2). In addition, there is a clear relationship between 24nt cross-talk siRNAs and methylation levels in the captured region of syntelogs (Table 1 and Table S3). This relationship is stronger for CHH methylation, which is more reliant on RdDM than CG and CHG methylation (3). The magnitude of the effect is not inconsequential, because the number of 24nt cross-talk siRNAs explains ~21% of CHH methylation variation across captured fragments. These results suggest that i) substantial RdDM activity occurs predominantly in the captured fragments of syntelogs, and that ii) this activity is likely driven by epigenetic cross-talk with the TEs that contain the captured sequences.

The conflict model further predicts that the epigenetic interactions should not proceed to the extent that gene expression is altered, because natural selection will conserve the function of important genes. We assessed function by comparing gene expression between donor and free syntelogs, and indeed found no evidence of reduction in expression of donor genes (Figure 2d, Figure S8); in fact, donor genes were more highly expressed than free genes. We propose that this difference likely reflects biases in capture events. This hypothesis presupposes that TEs are better able to capture highly expressed genes in open chromatin, and it conforms to the integration preferences of several TE families across plants and animals for genic regions (28, 44). Another interesting feature is the genic region that has been captured. Evidence suggests that methylation of the 5’ and 3’ UTRs - two regions with important regulatory roles for gene function (42, 43) - significantly affects expression levels in humans (45). Our analysis supports this claim by showing that capture of the 5’ or 3’ side of syntelog genes is associated with lower expression than capture of internal exons (Figure S9). Based on this negative effect, one expects 5’ and 3’ capture events to be rare for important genes or that natural selection quickly removes them from the population. However, we did not observe a scarcity of 5’ or 3’ captured fragments in donor syntelogs compared to donor translocated genes (17.3% vs. 15.6% for 5’ exons; 20.8% vs. 21.5% for 3’ exons); we hypothesize that their abundance might be linked to the time that it takes for siRNA crosstalk to establish.

Our results clearly illustrate the epigenetic effects of TE capture on donor syntelogs, but there are additional features that merit discussion. The first is the curious case of translocated genes (Figure 5b). Translocated genes are captured more often by TEs, because 16.5% of all translocated genes were found to be donors compared to only 5% of all syntelog genes. This pattern was especially evident in Helitrons (Figure S11a). Previous work has shown that TEs contribute to modifications of synteny (46), suggesting that TE capture can trigger gene movement. It is therefore possible that the categories of ‘donor’ and ‘translocated’ are linked mechanistically. Donor translocated genes are also highly methylated as most have >90% CG and CHG methylation (Figure 2c). This pattern is consistent with gene silencing, which is supported by siRNA mapping that is not specific to the captured region (Figures 3, S7), their very low expression levels (Figure 2d), and low percentage with functional annotation (64%). We propose that donor translocated genes are the exceptions that prove the rule - i.e., they illustrate the run-away effects of epigenetic interactions with TEs in the absence of selection for function. That said, it is worth emphasizing that the epigenetic patterns of donor translocated genes are not a general feature of translocated genes, because only a subset of free translocated genes has the combination of high methylation and low expression levels (Figure 2c,d). It is tempting to suggest that some genes of this subset represent donor genes for which we failed to identify their capture by a TE.

Another consideration is the case of directionality. Could it be that TEs simply tend to capture highly methylated genes? This notion can be rejected based on at least four pieces of information. First, most of the 1,629 donor genes are syntelogs (948, 58.2%) whose methylation levels are much lower compared to donor translocated genes. Second, this argument does not easily explain why the epigenetic effects are found only within the captured region of donor syntelogs. This difference is not likely to be a simple function of statistical power, because non-captured regions were consistently longer than captured regions. Third, the linear model (Table 1) establishes a positive relationship among capture, cross-talk siRNAs and methylation. Since siRNAs facilitate methylation via RdDM and not *vice versa*, this result implies a directionality that contradicts the simple explanation of a high methylation bias. Finally, we can use orthology relationships with sorghum to assess whether capture is biased toward genes that are highly methylated (Figure S12). We examined patterns of CG methylation between maize-sorghum syntelogs, separated between the free and donor categories based on our analysis in maize. Both free and donor genes show a positive correlation between species (Spearman coefficient r=0.62 for free and r=0.5 for donor genes, p<2.2e-16 for both), which reflects the well-established fact that genic methylation tends to be conserved over evolutionary time (24, 26, 47). More importantly, however, they imply that preexisting levels of methylation do not seem to trigger capture events, because genes located throughout the methylation spectrum have been captured in maize.

Lastly, we consider potential limitations to our analyses. We recognize, for example, that our set of donor genes does not represent all capture events throughout the history of the maize genome. This is because we did not examine all known TE families in maize and because we used criteria to identify capture events that were stricter than previous studies (e.g. *E*-value cutoffs of 1×10^-40^ vs. 1×10^-5^) (13, 14, 20, 23, 48), a conservative approach that favors specificity over sensitivity. As a result of these methodological decisions, our set of free genes must contain false negatives, i.e. undetected capture events. Similarly, our set of donor genes may also contain false positives. The crucial point about both false negatives and false positives is that they should reduce - and not enhance - epigenetic differences between donor and free genes. Hence, we suspect that, if anything, we have systematically underestimated the magnitude of epigenetic effects of TE capture on donor genes, at least for maize.

### The case for conflict: TEs

The conflict model also predicts that TEs with captured gene fragments gain an advantage. It is an open question as to how to measure such an advantage, and so we investigated several potential options. We asked, for example, whether TEs with fragments of syntelog genes have a tendency for camouflage, as measured by siRNA mapping or methylation levels. Consistent with the conflict model, these TEs map fewer siRNAs than free TEs, even when the captured region was masked (Figure 4b), and when TE age was taken into account. One caveat to this result is that we likely underestimated the size of the captured region; this could bias analyses if captured regions tend to map fewer siRNAs than TE-specific regions. TEs with syntelog capture also tend to have lower CG and CHG methylation than free TEs (Figure 4c). However, this is a nuanced result, for two reasons. First, we find that differences are small in magnitude and both TE types have >90% methylation on average. At these levels, any TE is probably effectively silenced. Second, TEs with syntelog capture events have ~3-fold higher levels of CHH methylation (Figure 4c), which is hard to reconcile with the lower number of matching siRNAs. The cause of this CHH difference remains elusive, but it contributes to the overall impression that these TEs have ongoing epigenetic interactions defined in large part by elevated CHH methylation levels for both the TE and the donor genes. Finally, if gene fragments provide camouflage for TEs, one reasonable prediction is that they will exist within the genome for longer periods of time than free TEs. We found that this is indeed the case (Figure 4a), but this result was principally caused by Sireviruses, perhaps in part reflecting their higher proportion of free TEs.

Altogether, we consider the case for TE advantage to be tantalizing and perhaps correct, but not yet fully convincing especially considering that we did not find evidence that these TEs reached high copy numbers after the capture events (Figure 4d). It is worth mentioning, however, an interesting alternative. It is possible that the epigenetic response against a TE continues unabated after the capture event, so that it gains no advantage, but the epigenetic effects on donor genes are moderated by natural selection using other mechanisms, such as active CHG demethylation (49). In such a scenario, further research will be needed to understand how demethylation only occurs in the subset of genes that are important for host function.

### Concluding remarks

The intragenomic conflict between TEs and host genomes described herein raises several questions for future investigation. For example, it is likely that our model applies generally to plant genomes because gene capture by TEs is a common occurrence (11, 12, 14, 15, 17, 50), but it remains to be seen if the conflict is more pervasive in species with higher methylation levels and TE load, as is often the case for large genomes (24, 25), or if it varies across TE types depending on their intrinsic transposition and capturing mechanisms. In addition, genes that have translocated from their syntenic loci account for a substantial proportion of the gene content of plants. To illustrate this, thousands of genes have lost synteny between maize inbred lines such as B73, Mo17, and W22 (39, 51). Further research is needed to show if capture by TEs is mechanistically linked with this gene movement; if true, then this path may represent a main route towards pseudogenization.

## Materials & Methods

### TE and gene datasets

For TEs, we utilized three published datasets that were carefully curated, representing full-length Helitrons, Pack-MULEs and Sireviruses. For Helitrons, we downloaded the coordinates on the B73 RefGen_v2 genome of 1,351 high-quality elements that were *in silico* validated with the Mo17 inbred line in Xiong et al. (2014). Reflecting their sequence quality, most of these elements have a local combinational value (LCV) score of >50. For Pack-MULEs, the coordinates of 275 full-length elements from Jiang et al. (2011) were based on the RefGen_v1 genome; hence, we aligned their sequences (BLASTN, *E*-value 1 x 10^-180^) on the RefGen_v2 genome requiring 100% identity on the complete length of each element. This approach yielded 251 Pack-MULEs. We note that the official RefGen_v4 TE annotation (B73v4.TE.filtered.gff3) contains 1,369 elements of the DNA Transposon Mutator (DTM) superfamily, some of which are presumed to be Pack-MULEs. This low number suggests that DTM and Pack-MULE elements are not abundant in the maize genome and that our dataset captures a substantial proportion of them. Finally, we downloaded from MASiVEdb (32) the sequences of 13,833 Sireviruses identified in RefGen_v2 using the MASiVE algorithm (52). We then filtered out elements that overlapped with each other and those with >5 consecutive ‘N’ nucleotides, based on evidence that BLASTN hits between genes and TEs often mapped precisely at the border of these stretches, indicating potential errors during scaffold assembly. To ensure that TEs are physically present in RefGen_v4, we converted their chromosomal coordinates from RefGen_v2 to RefGen_v4 using the Assembly Converter tool (http://www.gramene.org/) and only kept TEs with ≥90% of length converted on the same chromosome as RefGen_v2. We note that ~97% of the TEs that passed this filter had ≥99% length conversion. Our final TE population consisted of 1,035 Helitrons, 238 Pack-MULEs and 6,200 Sireviruses.

Our input for genes was the RefGen_v2 FGS (http://ftp.gramene.org/maizesequence.org/). We only included evidence-based genes and filtered for TE-related keywords using the annotation files ZmB73_5b_FGS_info.txt, ZmB73_5b_FGS.gff, ZmB73_5b_WGS_to_FGS.txt, ZmB73_5a_gene_descriptors.txt, ZmB73_5a_xref.txt. We also filtered for similarity (BLASTN, *E*-value 1 x 10^-20^) of the exons to the conserved domains of the reverse transcriptase and integrase genes of LTR retrotransposons using Hidden Markov Models (PF07727 and PF00665) from Pfam (https://pfam.xfam.org/). To remove genes that were no longer annotated in RefGen_v4, we linked the RefGen_v2 and RefGen_v4 gene IDs using files ‘updated_models’ in https://download.maizegdb.org/B73RefGenv3/ and ‘maize.v3TOv4.geneIDhistory.txt’ in http://ftp.gramene.org, and accessed information on function with ‘Zea_mays.B73_RefGen_v4.43.chr.gff3’ in http://ftp.gramene.org. These steps produced a final dataset of 27,056 genes. We finally assessed syntenic relationships with sorghum using data produced by Springer et al. (2018) and kindly provided to us by Dr. Margaret Woodhouse of MaizeGDB.

### Identification of capture events

We first removed all cases of physical overlaps between genes and TEs and then ran a BLASTN search between the exons of the longest transcript of each gene and our TE dataset. We opted for a strict *E*-value cutoff of 1×10^-40^, because we intended to minimize false positive events. The average capture length was 280nt, with a minimum of 90nt and a maximum of 1,932nt. When exons from multiple genes overlapped partially or fully with a TE, we selected the highest BLASTN bit score to define the true donor gene (20, 23). If exons from multiple genes had the same bit score, they were all regarded as true donors and kept for downstream analyses. Often, a TE contained multiple independent capture events, defined as non-overlapping areas within the TE. In total, we identified 6,838 such areas across all our TEs. We tested how this number changed after merging areas located in close proximity to each other, with the assumption that they may in reality represent a single capture event that BLASTN failed to identify in its entirety. By allowing a window of 10nt or 50nt, the number only slightly reduced to 6,724 and 6,379 respectively, suggesting that the majority represent truly independent capture events.

### siRNA, methylation and expression data

For siRNA mapping, we retrieved short read libraries for ear (GSM306487), leaf (GSM1342517) and tassel (GSM448857). We used Trimmomatic (53) to trim adaptor sequences, and FASTX toolkit (http://hannonlab.cshl.edu/fastxtoolkit/) to remove low quality nucleotides until reads had ≥3 consecutive nucleotides with a phred Q score >20 at the 3’ end. Reads of 21nt, 22nt and 24nt in length were kept and filtered for tRNAs (http://gtrnadb.ucsc.edu/), miRNAs (http://www.mirbase.org/), and rRNAs and snoRNAs (http://rfam.xfam.org/), and then mapped to the RefGen_v2 genome using BWA with default settings and no mismatches (54). Both uniquely and multiply mapping siRNAs were considered, and all loci of multiply mapping siRNAs were counted. We retrieved with a custom Perl script the number and IDs of all distinct siRNA sequences that mapped to a locus (e.g. captured region within the TE, or an exon) to calculate mapping of distinct siRNA sequences per kb as previously suggested (38). This metric collapses the number of reads in a library for a distinct sRNA sequence and it therefore permits the efficient calculation of the diversity and density of different siRNA sequences that map to a locus. Finally, using the siRNA IDs, we were able to identify siRNA cross-talk events.

For DNA methylation analysis, we used previously published BS-seq data from ear (SRA050144) and leaf (SRR850328). Reads were trimmed for quality and adapter sequences with Trimmomatic using default parameters and a minimum read length of 30nt (53). Trimmed reads were mapped to the RefGen_v2 genome using bowtie2 (v2.2.7, parameters: -N 0 -L 20 -p 2) within the bismark (v0.15.0) software suite (55). We did not allow mismatches and retained only uniquely mapped reads. The number of methylated and unmethylated reads at each cytosine in the genome was calculated using bismark_methylation_extractor. Positions with >2 reads were retained for further analysis. Bisulfite conversion error rates, or false methylation rates (FMR), were estimated from reads that mapped to the chloroplast genome. A binomial test incorporating the estimated FMR (*P*<0.05 after Benjamini-Yekutieli FDR correction) was then used to identify methylated cytosines (56). For each locus we retrieved the number of covered and methylated CG, CHG and CHH sites and calculated methylation levels for each context with ≥10 covered sites by dividing the number of methylated to covered cytosines (24). Methylation BS-seq data for the leaf tissue in sorghum were retrieved from reference (26).

Finally, we downloaded gene expression data from the ATLAS Expression database (www.ebi.ac.uk/gxa/) for ear (E-GEOD-50191), leaf (E-MTAB-4342) and various tissues of the maize kernel (E-GEOD-62778). Only genes with >0.1 TPM are included in the ATLAS database, hence we classified all other genes as having no expression.

### Statistical analyses of donor genes for siRNA mapping, expression and methylation

We used a one-sided binomial test to test whether cross-talk siRNAs map to donor genes more often than expected by chance. The number of successes is the number of cross-talk siRNAs, the number of trials is the total number of siRNAs that mapped to the donor gene, and the probability of success is the proportion of the total exonic length that has been captured by all TEs. If we assume a random distribution of siRNA across the donor gene, the expected probability of mapping of any siRNA to the captured area is the length of the captured area divided by total gene length. Binomial exact test p-values were corrected for multiple testing using Benjamini & Hochberg (1995).

In order to study the link between methylation levels of regions of donor genes and the number of cross-talk siRNAs, the lmer function of the R package lme4 (57) was used to write a linear model with mixed effects. The r.squaredGLMM function of the R package MuMIn (58) was used to compute the marginal R-squared (the variance explained by the fixed effects, here the number of cross-talk siRNAs). The proportion of methylated cytosines was log transformed, and the gene was set as a random factor (each gene had one measurement for leaf and one for ear). The analysis was repeated separately for the three methylation contexts and each siRNA length for captured and non-captured regions of donor genes:

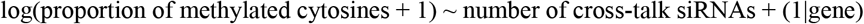

### Age of capture events

In order to estimate the age of gene capture events, we estimated synonymous divergence between donor genes and the captured fragments within TEs. The RefGen_v2 genome GFF file was used to split sequences into coding and non-coding (since in v2 UTRs are included in the first/last exons). The coding parts of donor genes and captured fragments were aligned using MACSE v2 (59). In cases where stop codons were found in the captured gene fragment, they were replaced by ‘NNN’ in order to compute synonymous divergence (dS) using the yn00 program in the paml package (60). To obtain capture age, dS values were divided by 2 x (1.3 x10-8) (61).

The lmer function of the R package lme4 (57) was used to write a generalized linear model with mixed effects to study the link between capture age and the number of cross-talk siRNAs. The r.squaredGLMM function of the R package MuMIn (58) was used to compute the marginal R-squared (the variance explained by the fixed effects, here capture age). The poisson family was used and gene was set as a random factor:

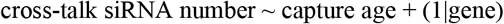

### Statistical analyses of TEs for siRNA mapping and methylation

For siRNA mapping, the glmer function of the R package lme4 (57) was used to write an exponential model with mixed effects to study the effect of TE type (with or without gene capture) and TE age. The r.squaredGLMM function of the R package MuMIn (58) was used to compute the marginal R-squared (the variance explained by the fixed effects, here TE type and age). The lsmeans function from the R package lsmeans (62) was used to compute the contrast between TEs with and without gene capture. The TE was set as a random factor (tissue was the repetition) and the number of siRNAs per kb was log transformed:

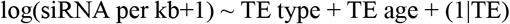

A simpler model was also used:

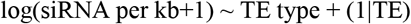

Similarly, a generalized linear model with mixed effects was used to study the effects of TE type and TE age on TE methylation. The binomial family was used, and TE was set as a random factor (tissue was the repetition). The analysis was repeated separately for the three methylation contexts:

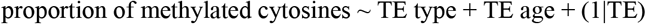

## Acknowledgements

A.M. is supported by an EMBO Postdoctoral Fellowship ALTF 775-2017 and by HFSPO Fellowship LT000496/2018-L. D.K.S. is supported by a Postdoctoral Fellowship from the National Science Foundation (NSF) Plant Genome Research Program (1609024). B.S.G. is supported by NSF grant 1655808. A.B. is supported by The Royal Society (award numbers UF160222 and RGF/R1/180006).

## Author Contributions

AB and BSG designed research; AM, ND, AB performed research; AM, DS, EP, AB analyzed data; AB and BSG wrote the paper.

