## Supplementary Figures for "Gene capture by transposable elements leads to epigenetic conflict in maize"

### Figure S1

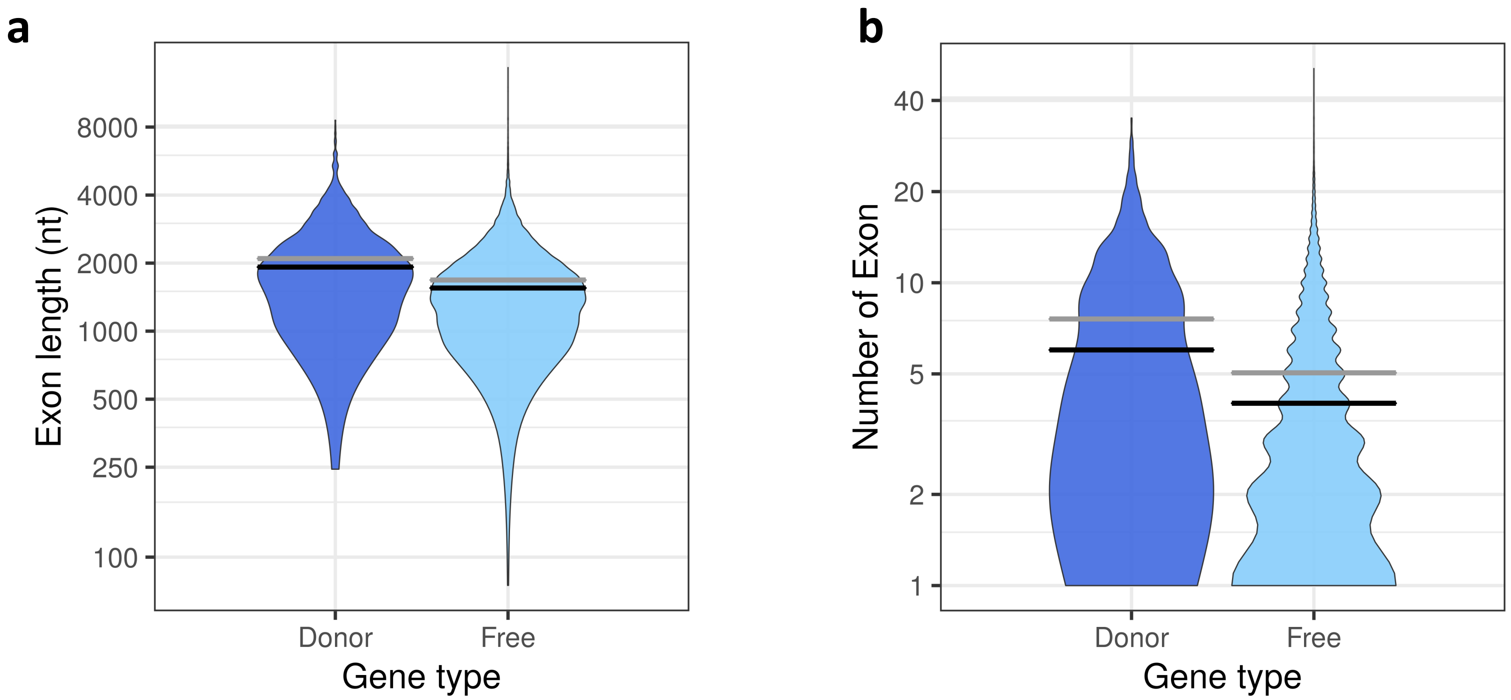


**Figure S1:** Comparison of the number of exons and length of the exonic sequence of donor and free genes. The gray lines indicate the mean, the black lines the median.

### Figure S2

**a**

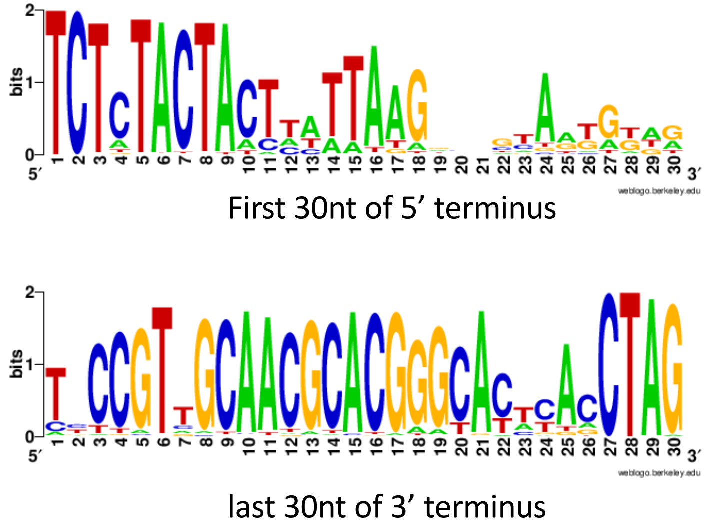


**b**

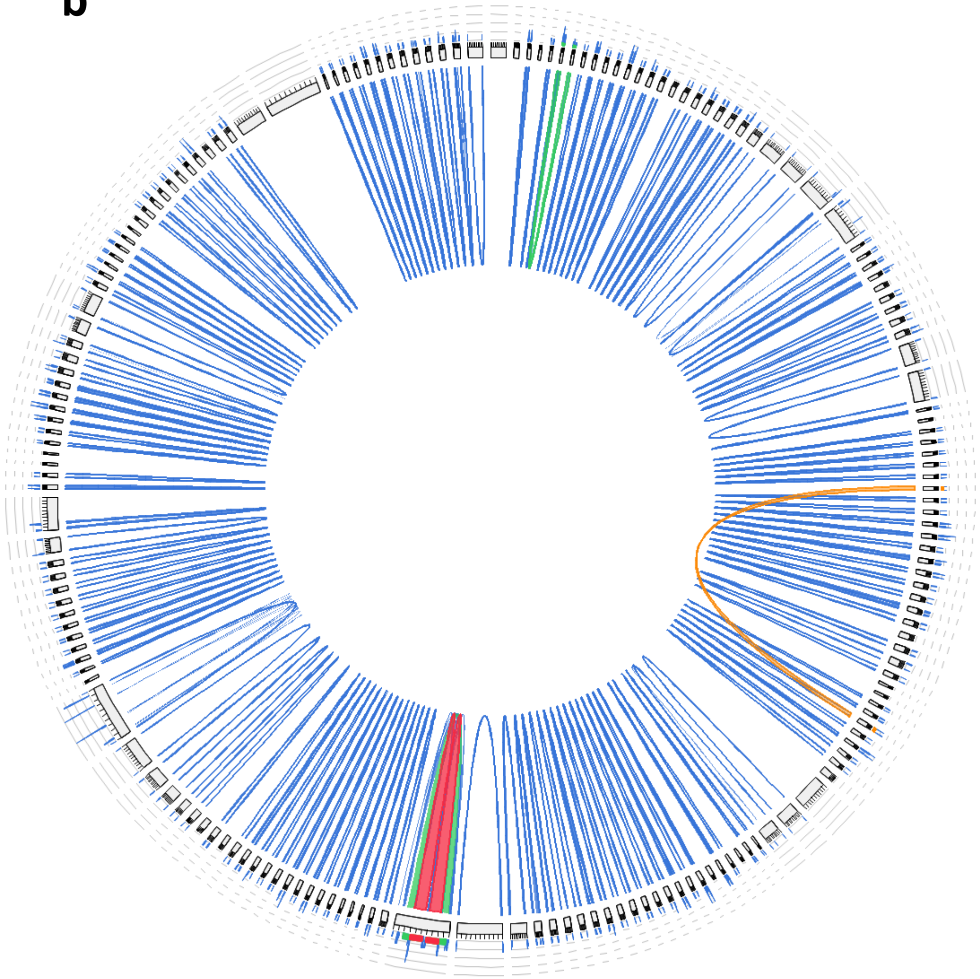

**c**

| TSD detected | Sireviruses |
|---|---|
| 5nt TSD | 164 |
| 4nt TSD | 29 |
| no TSD | 11 |
| total | 204 |

**Figure S2**. Structural and sequence features of full-length elements of the three TE families. **(a)** Helitron terminal conservation shown by sequence logos (https://weblogo.berkeley.edu/). Both the 5' and 3' ends are highly similar to those reported in the original HelitronsScanner manuscript by Xiong et al., 2014 (PMID: 24982153). **(b)** Conservation of the 5' and 3' terminal inverted repeats (TIRs) of Pack-MULEs. To calculate and visualize the sequence similarity of TIRs of individual elements we used Circoletto with a BLASTN E-value of $1x10^{-5}$ (Darzentas 2010, PMID: 20736339, http://tools.bat.infspire.org/circoletto/). Under this parameters, TIRs were detected for ~90% (167/186) of elements. **(c)** Using a custom Perl script, we searched for the typical 5nt target site duplication (TSD) generated by LTR retrotransposons upon integration. We detected a TSD for ~95% (193/204) of Sireviruses.

#### Figure S3

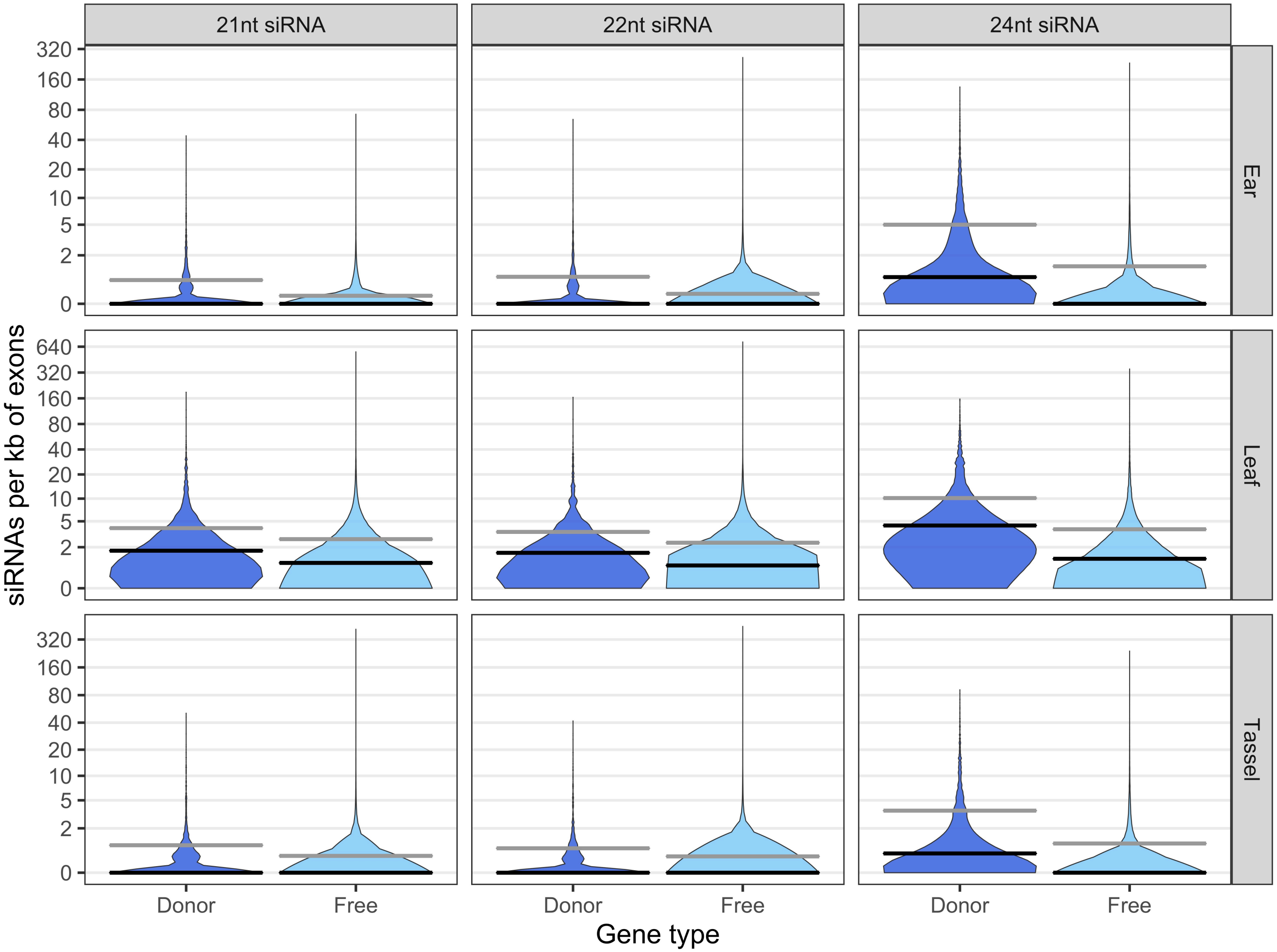


**Figure S3**. Number of 21nt, 22nt and 24nt distinct siRNA sequences per kb of exonic mapping to donor and free genes for ear, leaf and tassel tissues. Donor genes mapped significantly more siRNAs than free genes in all combinations (see main text for the statistical support). The gray lines indicate the mean, the black lines the median.

**Figure S4**

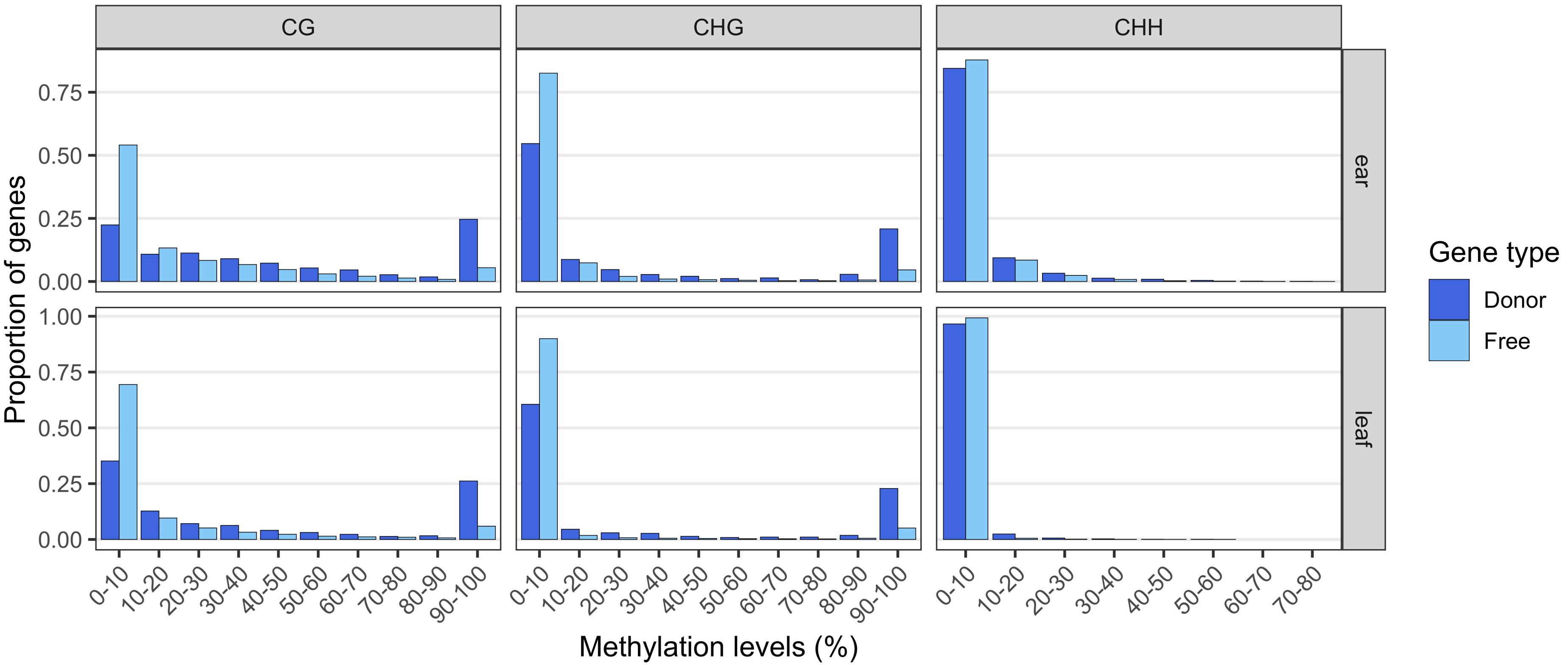


**Figure S4**. Distribution of the proportion of CG, CHG and CHH exonic methylation of donor and free genes in ear and leaf tissues. Donor genes were significantly more methylated in all combinations than free genes (see main text for the statistical support).

### Figure S5

**a**

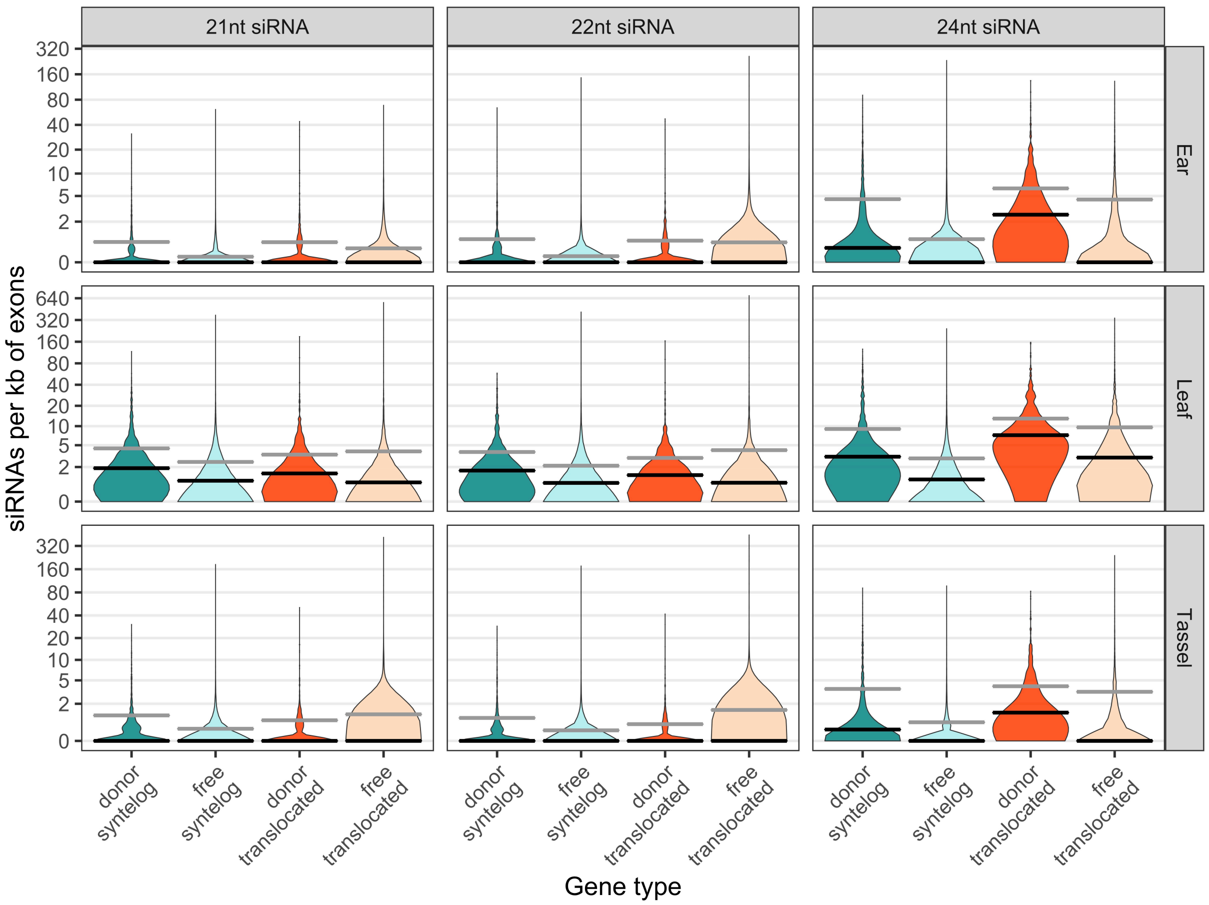


**b**

| one-sided Mann-Whitney U test p value (only genes with >0 siRNAs mapping) | | | |
|---|---|---|---|
| category | sRNA length | syntelogs | translocated |
| ear | 21 | 0.00105 | 0.9193 |
| | 22 | 3.79E-05 | 0.9999* |
| | 24 | 1.98E-12 | 0.1033 |
| leaf | 21 | 1.51E-05 | 0.4001 |
| | 22 | 9.10E-07 | 0.5706 |
| | 24 | 2.20E-16 | 6.89E-12 |
| tassel | 21 | 0.2489 | 0.9908* |
| | 22 | 0.06246 | 0.9996* |
| | 24 | 8.90E-14 | 0.3876 |

asterisks indicate cases were significantly more siRNA mapped to free than donor genes

**Figure S5**. siRNA profiles of donor and free genes split by their syntenic status with sorghum for ear, leaf and tassel tissues. The four gene categories in all plots are donor syntelogs, free syntelogs, donor translocated and free translocated genes. **(a)** Number of 21nt, 22nt and 24nt distinct siRNA sequences per kb of exonic mapping. Donor genes mapped significantly more siRNAs than free genes in all combinations (see main text for the statistical support). The gray lines indicate the mean, the black lines the median. **(b)** P values of a one-sided Mann-Whitney U test for siRNA patterns as in (a) but after removing genes with no mapped siRNAs. Patterns did not change for syntelogs, but did so for translocated genes because donor and free genes were equally mapped by 21-22nt siRNAs, or donor genes were significantly less mapped than free genes for 24nt siRNAs.

### Figure S6

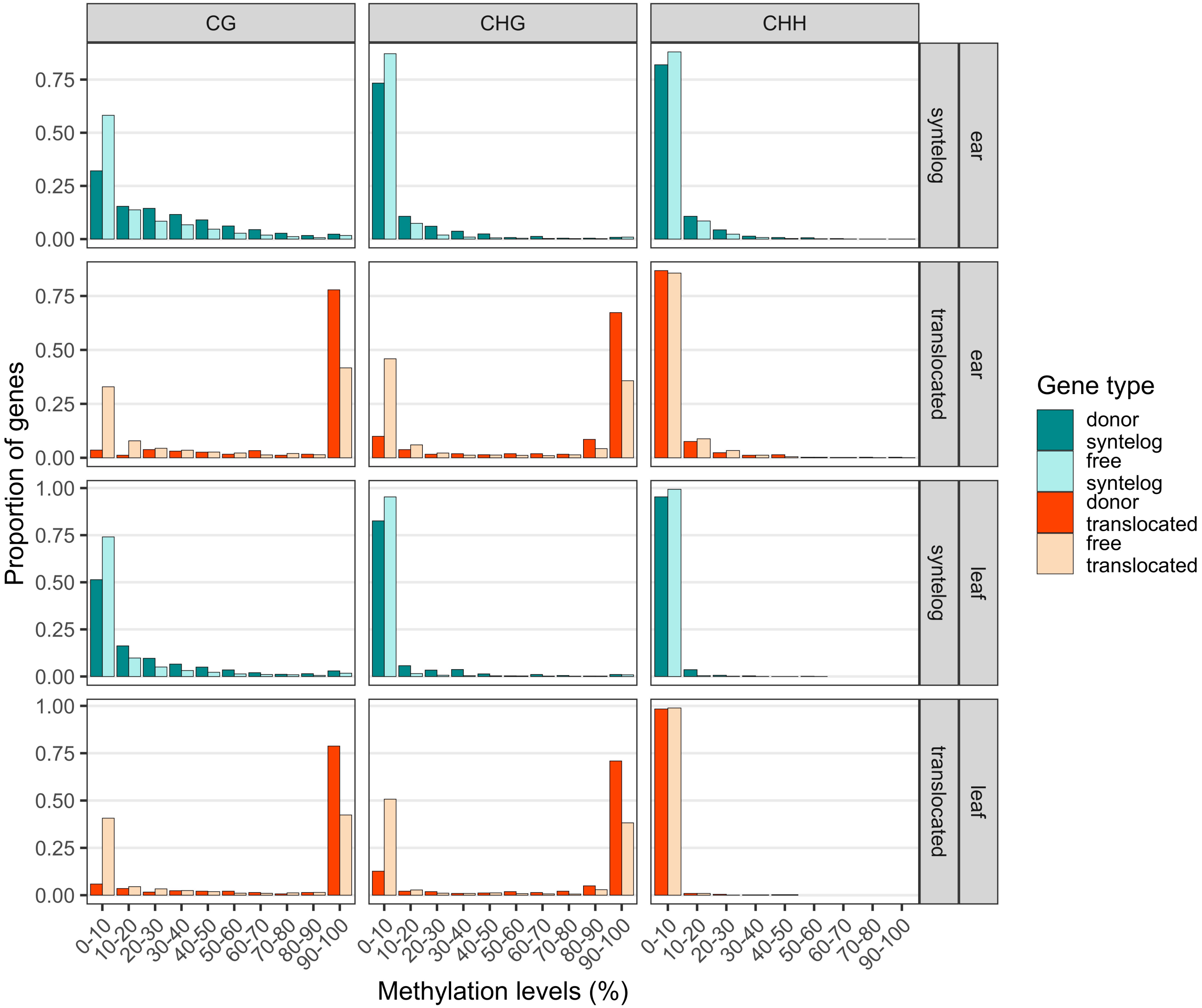


**Figure S6**. Distribution of the proportion of CG, CHG and CHH exonic methylation of donor and free genes split by their syntenic status with sorghum for ear and leaf tissues. The four gene categories in all plots are donor syntelogs, free syntelogs, donor translocated and free translocated genes. Donor genes were more methylated than free genes in all methylation contexts (see main text for the statistical support).

### Figure S7

**a**

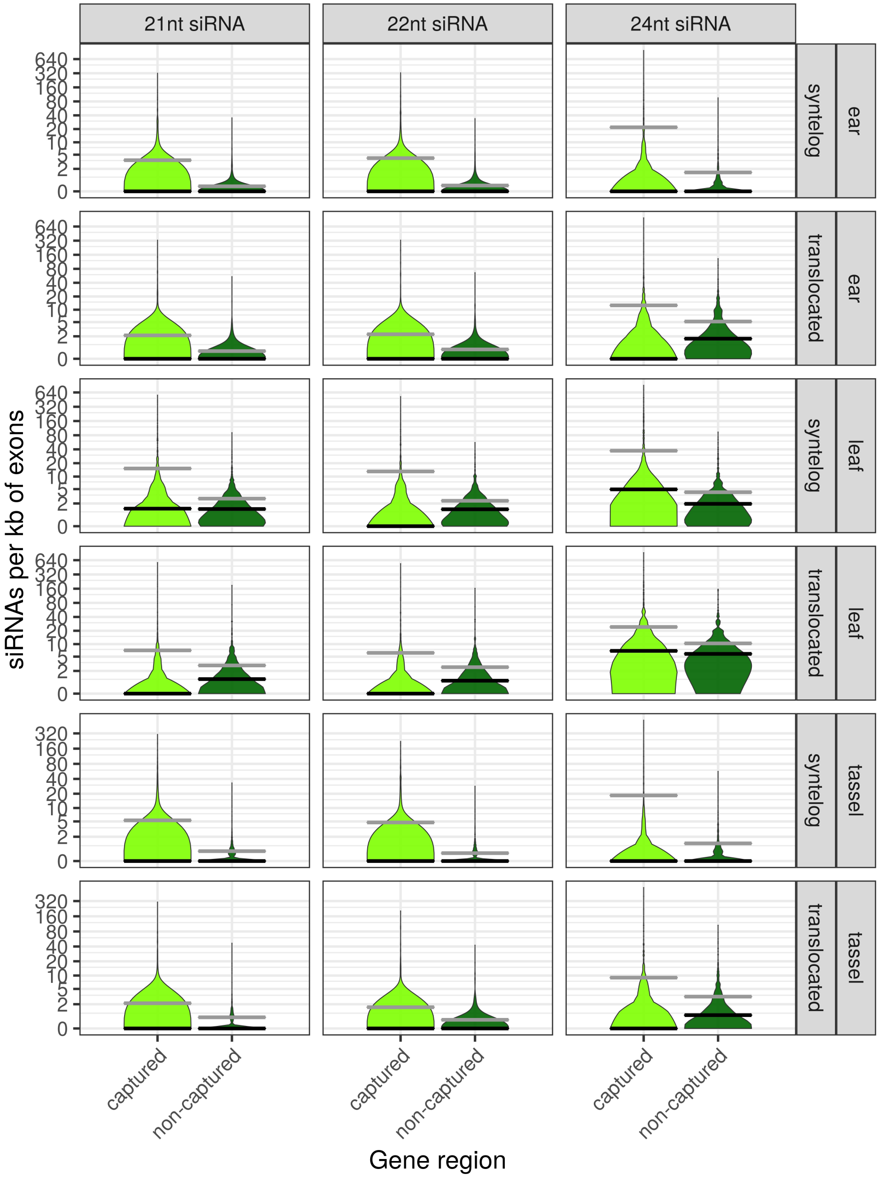


| one-sided Wilcoxon signed rank test p value for siRNA mapping of captured vs. non-captured regions | | | |
|---|---|---|---|
| tissue | sRNA length | syntelogs | translocated |
| ear | 21 | 2.20E-16 | 0.1536 |
| | 22 | 2.20E-16 | 0.3717 |
| | 24 | 2.20E-16 | 0.4025 |
| leaf | 21 | 4.26E-16 | 0.4322 |
| | 22 | 4.62E-10 | 0.4159 |
| | 24 | 2.20E-16 | 0.0005942 |
| tassel | 21 | 1.71E-11 | 0.5653 |
| | 22 | 1.32E-08 | 0.1247 |
| | 24 | 2.20E-16 | 0.003063 |

**Figure S7**. siRNA and methylation patterns of captured and non-captured regions of syntelog and translocated donor genes for ear, leaf and tassel tissues. **(a)** Number of 21nt, 22nt and 24nt distinct siRNA sequences per kb of captured and non-captured exonic regions. The gray lines indicate the mean, the black lines the median. **(b)** Distribution of the proportion of CG, CHG and CHH exonic methylation of captured and non-captured regions. The tables on the right include the p values of a one-sided Wilcoxon signed rank test for each combination shown in the plots.

### Figure S7 (cont.)

**b**

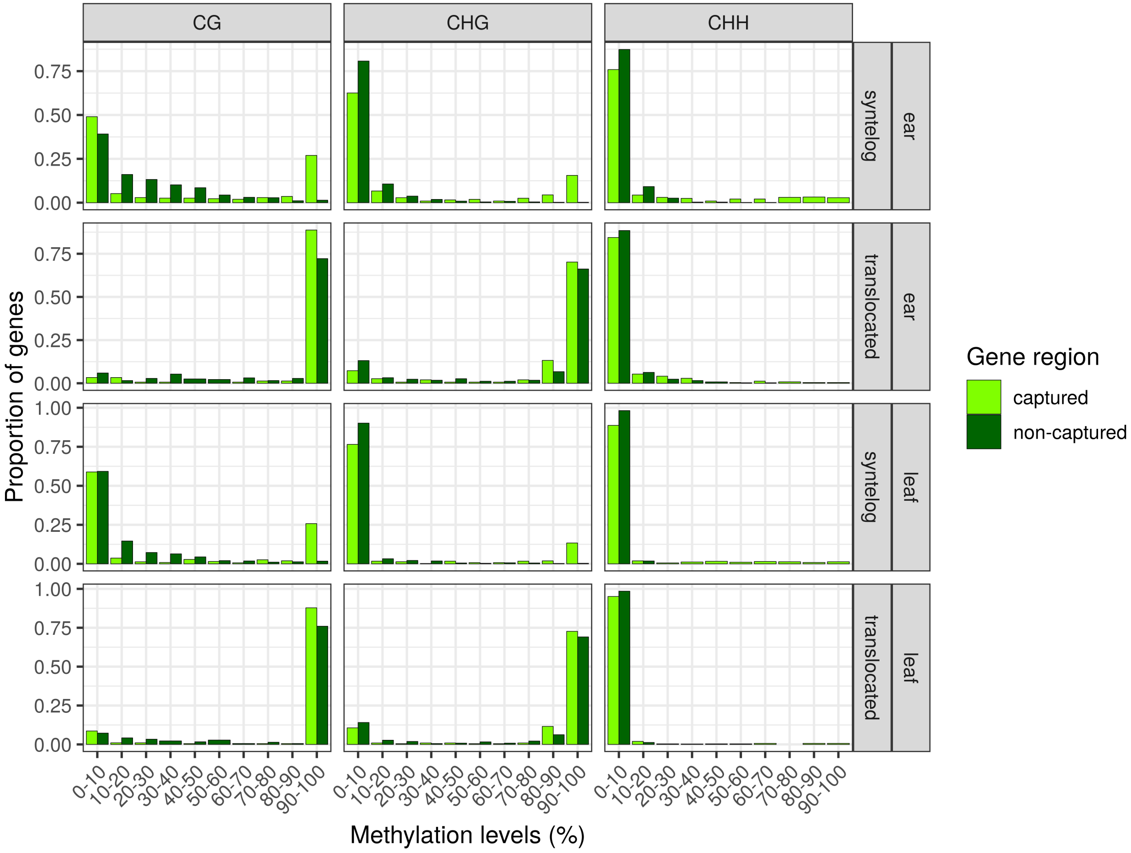


| one sided Wilcoxon signed rank test<br>p value for methylation levels of captured vs. non-captured regions | | | |
|---|---|---|---|
| tissue | methylation context | syntelogs | translocated |
| ear | CG | 9.45E-12 | 0.01012 |
| | CHG | 4.36E-12 | 0.08452 |
| | CHH | 1.49E-05 | 0.4433 |
| leaf | CG | 2.20E-16 | 0.04854 |
| | CHG | 2.20E-16 | 0.2596 |
| | CHH | 1.17E-10 | 0.2332 |

### Figure S8

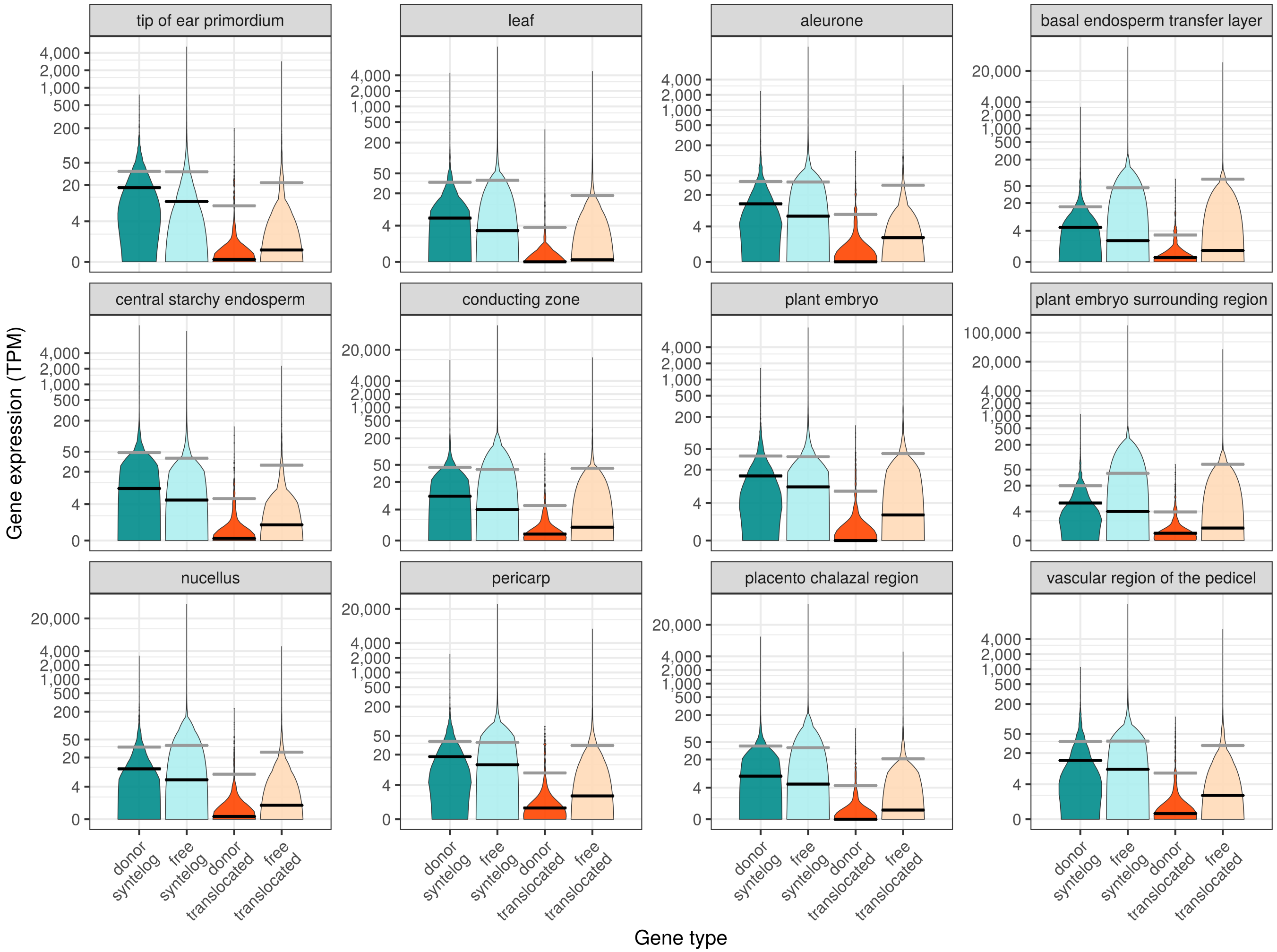


**Figure S8**. Expression profiles of donor and free genes split by their syntenic status with sorghum across different tissues. The four gene categories are donor syntelogs, free syntelogs, donor translocated and free translocated genes. Gene expression was measured in transcripts per million (TPM). Donor syntelogs were expressed at significantly higher levels than free syntelogs (one sided Mann-Whitney U test p-value in tip of ear primordium p<2.2e-16, leaf p=1.372e-15, aleurone p=2.346e-16, basal endosperm transfer layer p=2.518e-16, central starchy endosperm p=7.746e-14, conducting zone p=4.119e-16, plant embryo p=2.37e-15, plant embryo surrounding region p=1.041e-15, nucellus p=2.441e-13, pericarp p=3.85e-13, placento-chalazal region p=2.685e-08, vascular region of the pedicel p=3.044e-10), and donor translocated genes were expressed at significantly lower levels than free translocated genes (one sided Mann-Whitney U test p-value in tip of ear primordium p=3.129e-05, leaf p=8.232e-12, aleurone p=4.546e-08, basal endosperm transfer layer p=3.227e-06, central starchy endosperm p=1.291e-06, conducting zone p=7.005e-06, plant embryo p=4.304e-07, plant embryo surrounding region p=1.514e-05, nucellus p=0.000959, pericarp p=1.765e-06, placento-chalazal region p=6.866e-05, vascular region of the pedicel p=9.155e-07). The gray lines indicate the mean, the black lines the median.

#### Figure S9

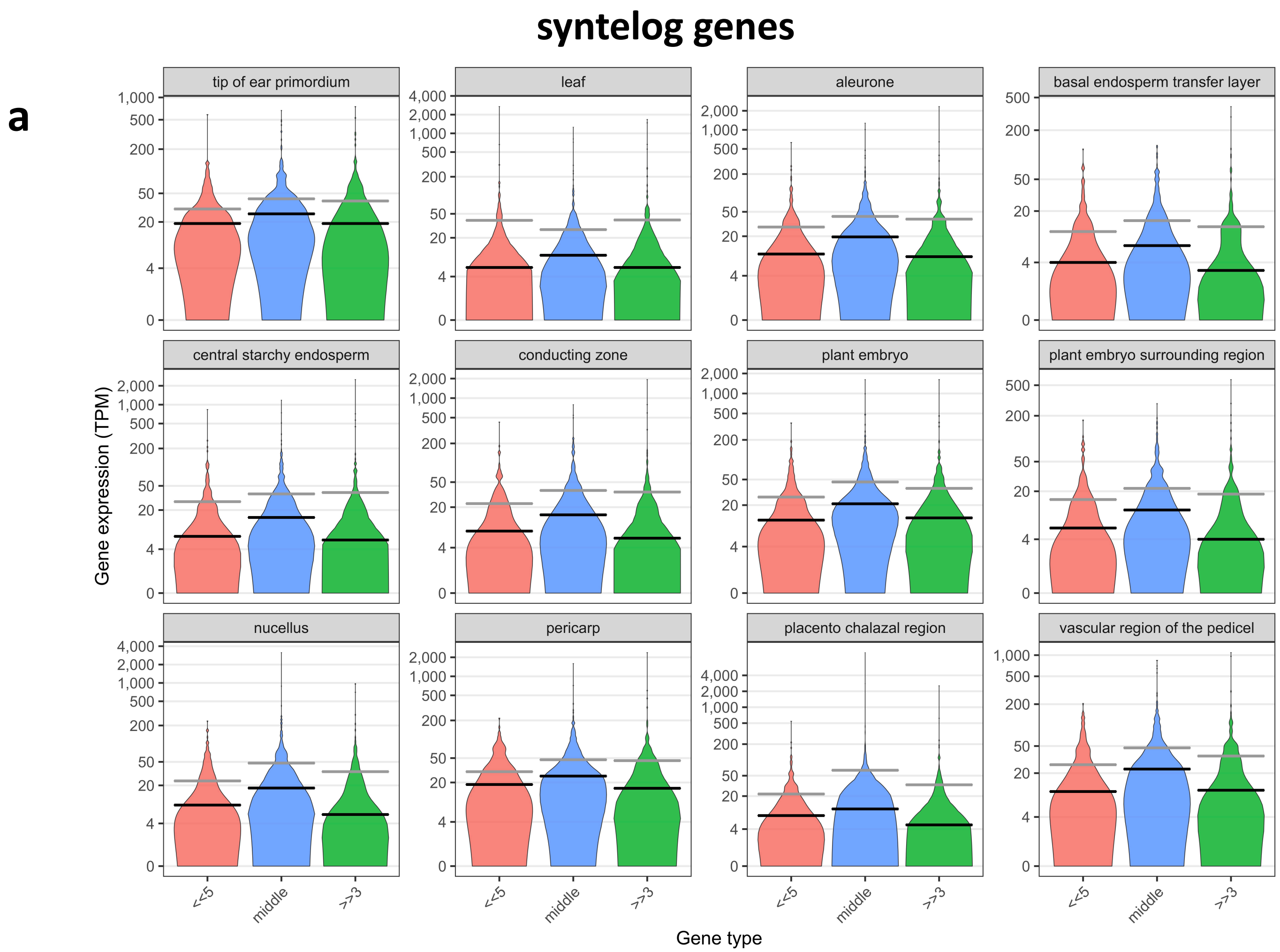


**Figure S9.** Expression profiles of donor syntelog and donor translocated genes split by the position of the captured fragment. The three position categories are genes with captured fragments only from the 5' exon, from the 3' exon, or any internal exon. Gene expression was measured in transcripts per million (TPM). (**a**) For syntelogs, capture of the 5' or 3' exons significantly reduced expression compared to internal exons (one-sided Mann-Whitney U test p-value in tip of ear primordium p=0.002 and p=0.005929, leaf p=0.01084 for 5' vs. internal exons and p=0.01399 for 3' vs. internal exons, aleurone p=3.442e-05 and p=4.303e-08, basal endosperm transfer layer p=0.000632 and p=1.066e-06, central starchy endosperm p=0.0004491 and p=1.305e-06, conducting zone p=0.0003493 and p=1.247e-06, plant embryo p=2.045e-05 and p=3.411e-06, plant embryo surrounding region p=0.0001193 and p=1.334e-06, nucellus p=0.0002257 and p=4.239e-07, pericarp p=0.004168 and p=7.079e-05, placento-chalazal region p=0.007731 and p=1.392e-05, vascular region of the pedicel p=0.0001092 and p=2.667e-05). (**b**) For translocated genes, there was no significant difference in expression between 5' or 3' exons vs. internal exons (tip of ear primordium p=0.2787 and p=0.8832, leaf p=0.5292 and p=0.5701, aleurone p=0.7191 and p=0.9771, basal endosperm transfer layer p=0.8982 and p=0.9509, central starchy endosperm p=0.6854 and p=0.8472, conducting zone p=0.8194 and p=0.957, plant embryo p=0.5082 and p=0.7896, plant embryo surrounding region p=0.7891 and p=0.9984, nucellus p=0.4451 and p=0.6221, pericarp p=0.7388 and p=0.9473, placento-chalazal region p=0.6112 and p=0.9378, vascular region of the pedicel p=0.4224 and p=0.975). The gray lines indicate the mean, the black lines the median.

#### Figure S9 (cont.)

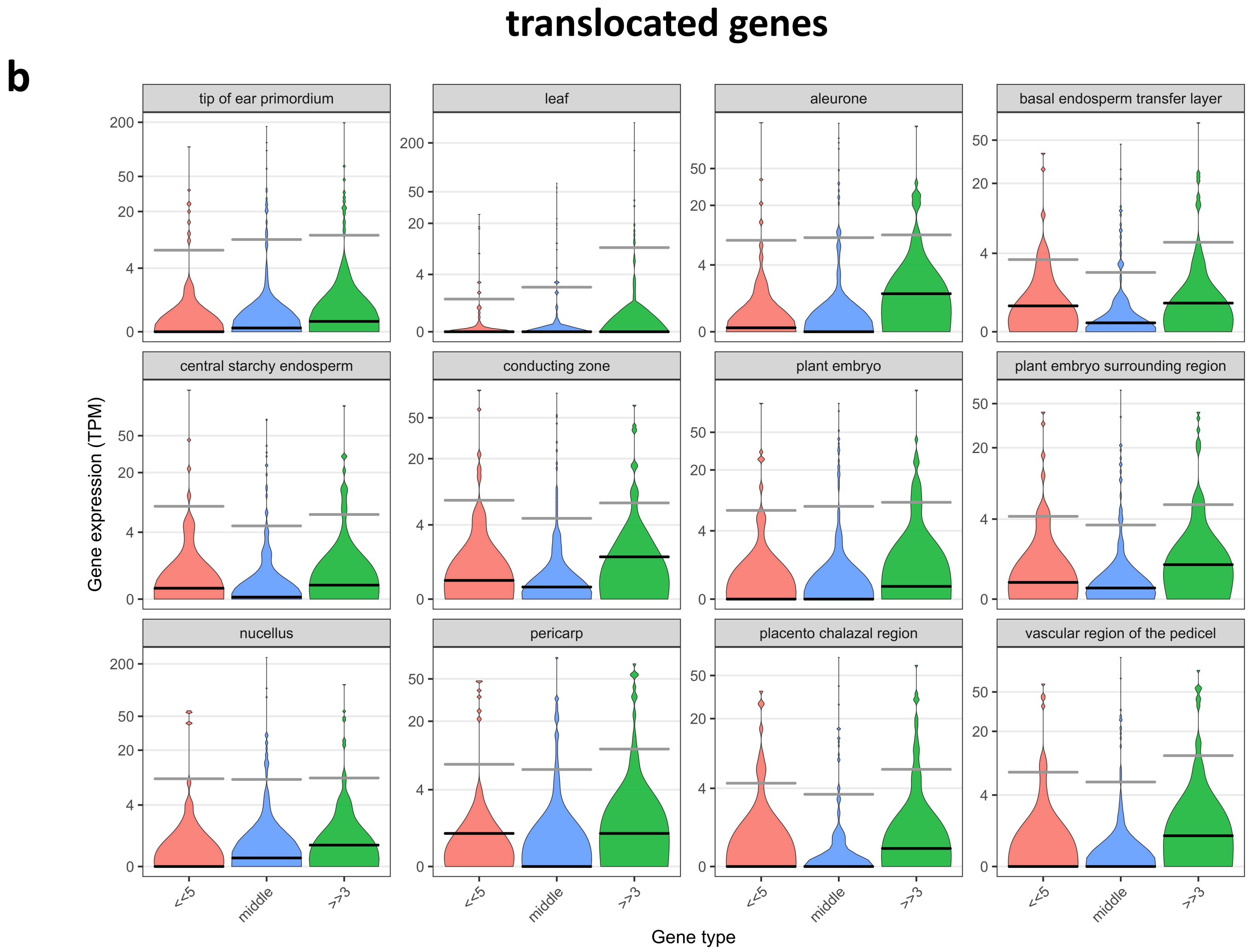

### Figure S10

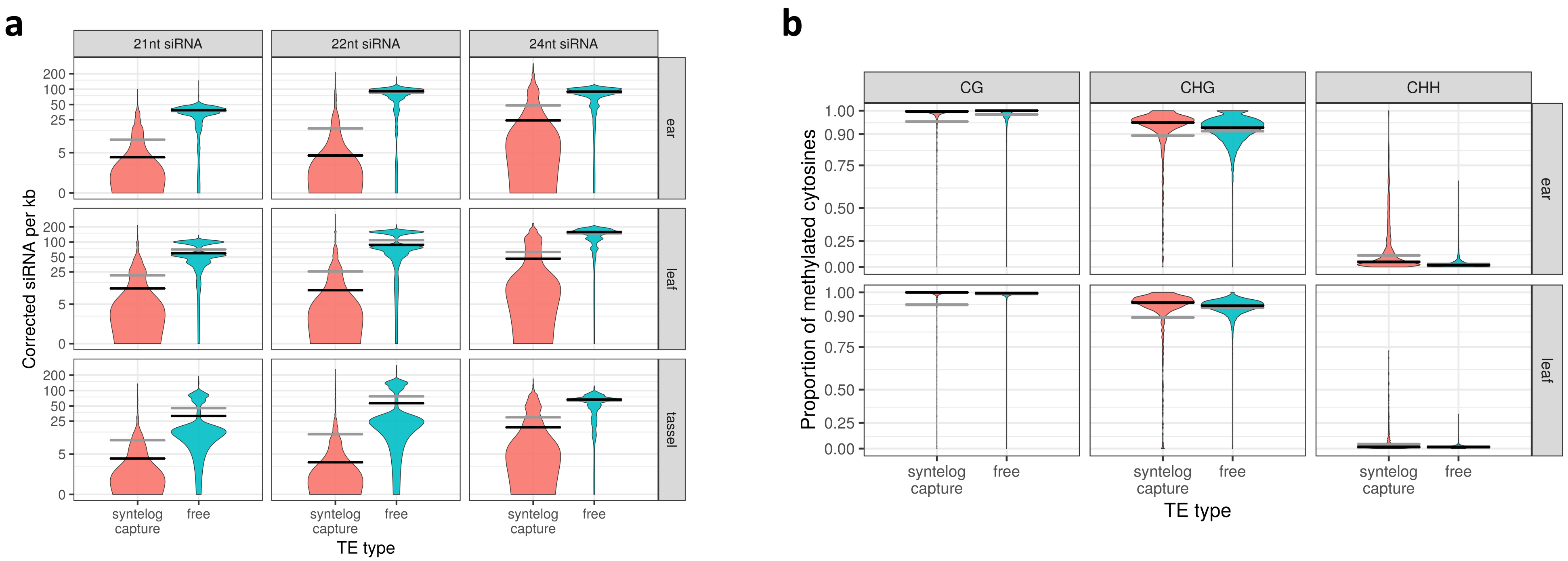


**Figure S10**. siRNA and methylation patterns of TEs with syntelog capture vs. free TEs for ear, leaf and tassel tissues. **(a)** Number of 21nt, 22nt and 24nt distinct siRNA sequences per kb mapping to TEs. This was computed after removing captured regions from TEs, but results were qualitatively identical when they were included. **(b)** Proportion of methylated cytosines in CG, CHG and CHH contexts of TEs. The gray lines indicate the mean, the black lines the median.

### Figure S11

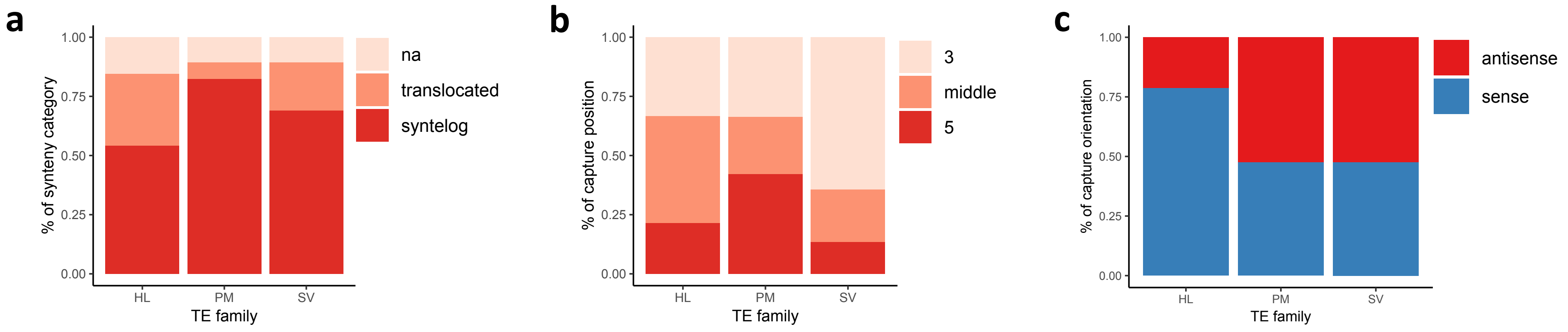


**Figure S11**. Capture characteristics of Helitrons, Pack-MULEs and Sireviruses. **(a)** Syntenic status of genes. The 'na' category includes both genes that were located in regions of maize chromosomes that were not identified in sorghum and genes for which no information was found regarding their synteny. **(b)** Position of the captured fragment within the gene sequence. This was allocated as 5' exon, 3' exon or any other internal exon. Genes that consisted of only one or two exons were excluded. (**c**) Orientation of the captured fragment in relation to the TE. HL, Helitrons; PM, Pack-MULEs; SV, Sireviruses.

### Figure S12

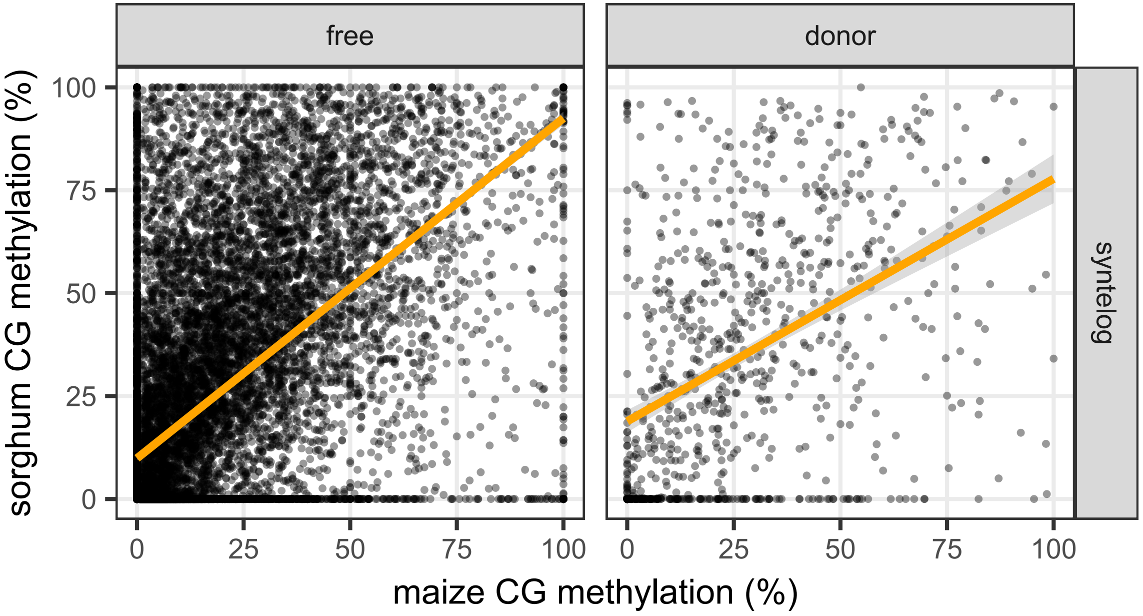


| Spearman coefficient r | |
|---|---|
| free | r = 0.62<br>p <2e-16 |
| donor | r = 0.5<br>p <2e-16 |

**Figure S12**. Relationship between CG methylation levels of syntelog genes of maize and sorghum. Genes were split in two groups based on if they were captured by a TE in maize. Each dot represents a pair of syntelogs. We identified the syntelog pairs using data from Springer et al., (2018, PMID 30061736) as described in the main text. We required that both genes of a pair had ≥10 covered CG sites (899 pairs for donor syntelogs and 16578 pairs for free syntelogs passed the filter). The shading around each regression line represents the 0.95 confidence interval, with the Spearman coefficient r and p value shown as a separate table on the right.
